# The NanoBridge system for targeted post-translational modification of challenging proteins

**DOI:** 10.64898/2026.06.08.730974

**Authors:** Fangfang Shen, Olga D. Merino-Chavez, Sheng-Yao Dai, Sebastian Alfonso, Laura M. K. Dassama

## Abstract

The ability to edit posttranslational modifications (PTMs) of endogenous proteins within cells is essential for precisely delineating the biological roles of PTMs and for developing targeted therapeutics. While the paradigm of chemically induced proximity (CIP) has advanced this field by enabling the recruitment of PTM enzymes to the proximity of proteins of interest (POIs), CIP requires small-molecule binders that are difficult to obtain for proteins without well-defined binding pockets. In principle, the use of biomolecular ligands that target disordered proteins should overcome this limitation. In this work, we developed the NanoBridge as a modular and generalizable platform to enable PTM editing of challenging POIs in live cells. The NanoBridge employs biologic binders to transiently direct the small protein tag FKBP12^F36V^ to unmodified target proteins, thereby enabling multiplex PTM editing upon use of heterobifunctional small molecules that recruit endogenous PTM enzymes. Compared with existing approaches, the NanoBridge offers greater flexibility to induce multiple types of PTMs on the same POI while providing precise temporal control and avoiding the introduction of exogenous PTM writers. Using eight protein binders targeting three structurally diverse and largely unstructured proteins - BCL11A (a hemoglobin regulator), KRAS (a cancer driver), and p53 (a tumor suppressor) – the NanoBridge mediated targeted degradation, phosphorylation, and acetylation in a rapid, reversible, and temporally controlled manner. As such, the NanoBridge represents a versatile strategy for the targeted modulation of endogenous proteins, particularly those lacking accessible small molecule ligands, and presents new opportunities for investigating the physiology of PTMs on challenging proteins.

## Introduction

Posttranslational modifications (PTMs) diversify protein function through covalent modification of amino acid residues following protein translation.^1^ The number of human protein isoforms generated through gene expression has been estimated to be around 100,000, which pushes the number of total proteins to the tens of millions after the consideration of splicing variants and PTMs. To date, more than 400 types of PTMs have been identified,^2^ and these modifications regulate protein conformation^3^, localization, interactions^4^, and signaling pathways^5,6^. Despite their critical roles, elucidating PTM function remains challenging because PTMs are often transient, low in abundance, and engage in extensive crosstalk that dynamically regulates protein function. Traditional methods that perturb a single PTM enzyme or manipulate a specific site are therefore limited in capturing this combinatorial complexity.

Inspired by the success of proteolysis targeting chimeras (PROTACs) and molecular glue degraders for targeted protein degradation,^7–10^ chemically-induced proximity (CIP) has now expanded to induce other protein modifications in ways that modulate the function of target proteins. By bringing endogenous PTM writers or erasers into proximity of the POI, ^11,12^ diverse PTMs can be induced or removed, even on POIs that are not natural substrates of the PTM enzymes. These emerging CIP tools include deubiquitinase-targeting chimeras (DUBTACs),^13^ acetylation tagging system (AceTAG),^14,15^ phosphorylation targeting chimeras (PhosTACs),^16^ phosphatase recruitment chimeras (PHORCs),^17^ and phosphorylation-inducing chimeric small molecules (PHICS),^18,19^ which all enable the targeted installation or removal of PTMs in living cells with improved temporal precision and substrate selectivity. However, extending CIP-based PTM editing to a broader range of substrates, particularly intrinsically disordered proteins (IDPs), remains challenging. Because CIP modulators are small molecules and are limited to proteins with well-defined pockets, their targets exclude many disease-relevant proteins that are IDPs whose functions are linked to the PTMs they carry.^20^ To overcome this limitation, the field has used genetically encoded tags fused to POIs^21–26^ to enable recruitment of PTM enzymes through tag-directed ligands (**Figure 1a**).^14–16,18,27^ While useful, an important drawback is that the introduction of fusion tags via gene modification may alter the native folding or cellular function of POIs. An alternative approach is to employ biologic binders to create binder–enzyme fusion systems, where the biologic is directly fused to the PTM modifier. While most of these systems have focused on targeted protein degradation,^28,29^ a few exciting studies have demonstrated the feasibility of using biologic binder–enzyme fusions to modulate targeted PTMs, such as glycosylation^30,31^ or phosphorylation^32^. However, these direct fusion strategies require the introduction of exogenous PTM enzymes into cells, potentially perturbing endogenous PTM homeostasis and generating non-physiological modifications. Therefore, methods capable of recruiting endogenous PTM machinery to endogenous IDPs could provide powerful opportunities for elucidating PTM functions and advancing therapeutic development.

**Figure 1.**
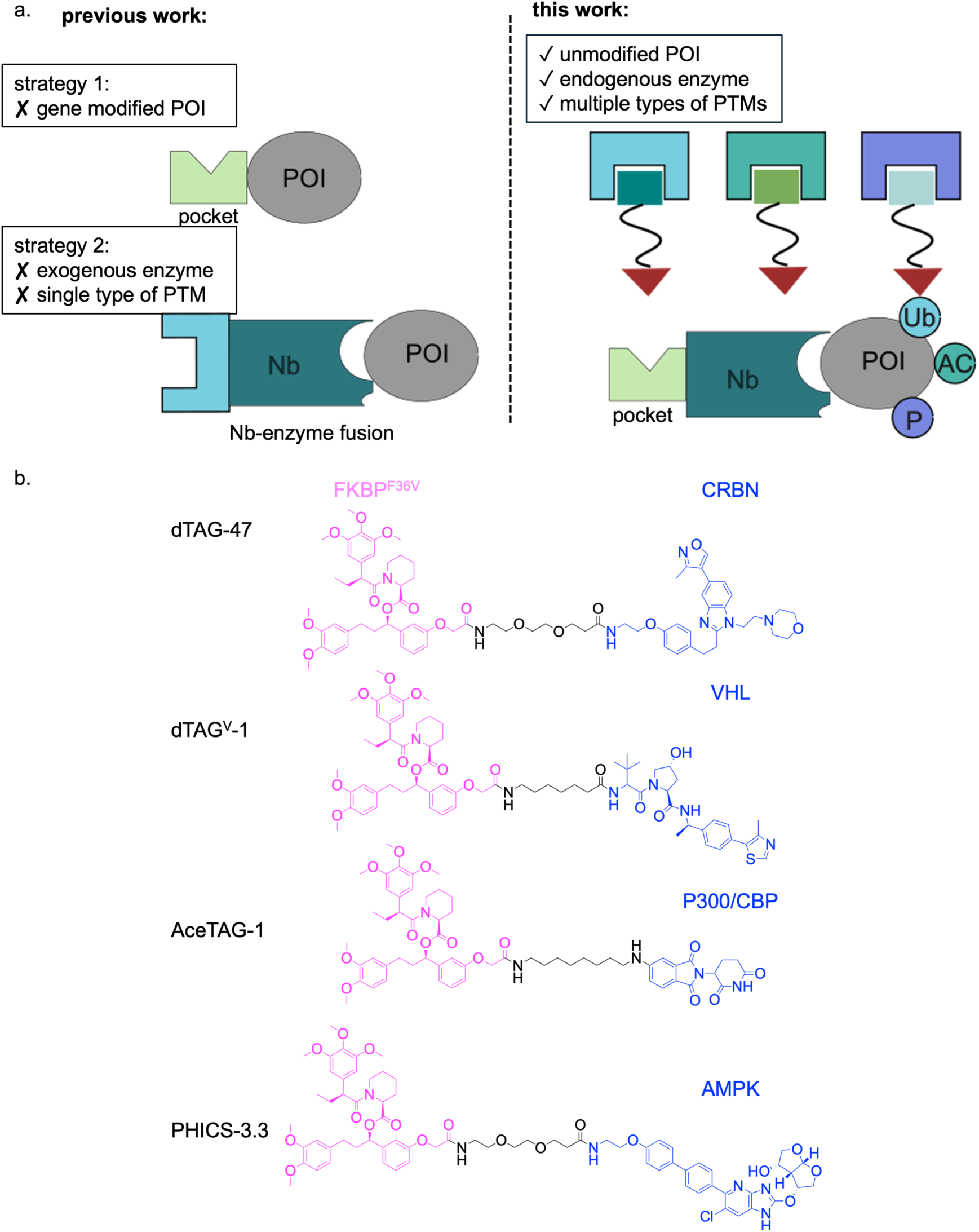
(a) Schematic overview of existing technologies *vs* the NanoBridge. Current technologies to access challenging targets use genetic editing to insert exogenous tags into N- or C-terminus of the POI or employ exogenous binder-enzyme fusions to target POIs. The NanoBridge instead uses a POI binder to direct a transient tag to various positions of POI. (b) Structures of the heterobifunctional ligands used in this study to induce PTMs on POIs.

In this work, we report the NanoBridge system that relies on biologic binders fused to the FKBP12^F36V^ tag to transiently direct ligandable pockets to IDPs. By leveraging previously reported heterobifunctional small molecules that bind FKBP12^F36V^ and PTM enzymes, we induced multiple types of PTMs on genetically unmodified proteins (**Figure 1a**). We demonstrated that NanoBridge is modular and compatible with biologic binders that include nanobodies and DARPins spanning diverse epitopes and affinities against different POIs. Unlike genetic tag fusion strategies, biologic binding is reversible and allows targeting POIs beyond the N- or C-terminal insertion sites typically used for genetic tags. In addition, the use of small molecules enables the recruitment of multiple endogenous PTM enzymes with temporal and dose-dependent control, which provides greater flexibility than direct fusion approaches. By simply switching proximity-inducing heterobifunctional small molecules, the NanoBridge induced protein degradation, acetylation, and phosphorylation of untagged BCL11A, KRAS, and p53 in living cells. As additional CIP strategies emerge, the NanoBridge could provide a versatile platform for studying and regulating PTMs on endogenous proteins, especially those that lack accessible small-molecule ligands.

## Results

### NanoBridge induced targeted degradation of BCL11A

BCL11A is a transcription factor whose downregulation is known to reactivate fetal-type hemoglobin.^33–36^ Casgevy, a recently approved therapy, utilizes the CRISPR/Cas9 system to edit the BCL11A enhancer for the treatment of sickle cell disease (SCD) and 𝛽-thalassemia. However, the high cost of gene therapy limits its broad application. TPD provides an alternative way to deplete BCL11A in a temporal, reversible, and cost-effective manner. Using yeast surface display technology, we identified specific nanobodies capable of targeting zinc finger domains (ZnFs) or their surrounding regions in BCL11A.^29,37^ In prior work, we fused a BCL11A specific nanobody to a cell permeant miniature protein and the catalytic domain of an E3 ligase to create cell permeant degraders of the highly disordered transcription factor^29^. However, this degrader lacked temporal control for the degradation, and the nanobody fusion was limited to degradation only. To evaluate the feasibility of the NanoBridge system for mediating BCL11A degradation in a temporal fashion, we fused nanobody 6101.19 (Nb6101.19),^37^ which binds to the ZnF456 domain of BCL11A with an equilibrium dissociation constant (*K*_d_) of 21.9 ± 0.4 nM, to FKBP12^F36V^. With this design, we expect a transient and reversible attachment of FKBP12^F36V^ to BCL11A through the binding of Nb6101.19. Using dTAG-47, a heterobifunctional molecule binding to FKBP12^F36V^ and the E3 ligase cereblon (CRBN), we aimed to induce the targeted degradation of BCL11A (**Figure 2a**). Human umbilical cord blood-derived erythroid progenitor cells (HUDEP-2) stably expressing strep-Nb6101.19-FKBP12^F36V^-NLS (to enhance nuclear localization) were obtained by lentiviral transduction, followed by puromycin selection. The addition of varying concentrations of dTAG-47 to these cells for 24 h led to a dose-dependent degradation of BCL11A, with maximum degradation achieved between 150 nM and 1.2 μM of dTAG-47 added (**Figures 2b, c**); the NanoBridge construct Nb6101.19-FKBP12^F36V^ was also degraded upon dTAG-47 treatment. Notably, the so-called “hook effect” was observed when dTAG-47 concentrations were higher than 600 nM, hinting that the observed degradation of NanoBridge is a consequence of ternary complex formation between dTAG-47, the NanoBridge, and CRBN. We next performed a time course experiment to monitor the kinetics of BCL11A and the NanoBridge. Although NanoBridge was degraded earlier than BCL11A, the remaining level of NanoBridge was sufficient to induce BCL11A degradation (**Figures 2d, e**). BCL11A loss is apparent after 8 hours of exposure to dTAG-47 (**Figures 2d, 2e**) and continues to decline for 72 h (**Figure 2f**). Additionally, dTAG-47 removal from cells results in a steady recovery of BCL11A to pre-treatment levels within 24 h (**Figure 2f**), indicating that the degradation is reversible and dependent on the presence of the heterobifunctional molecule.

**Figure 2.**
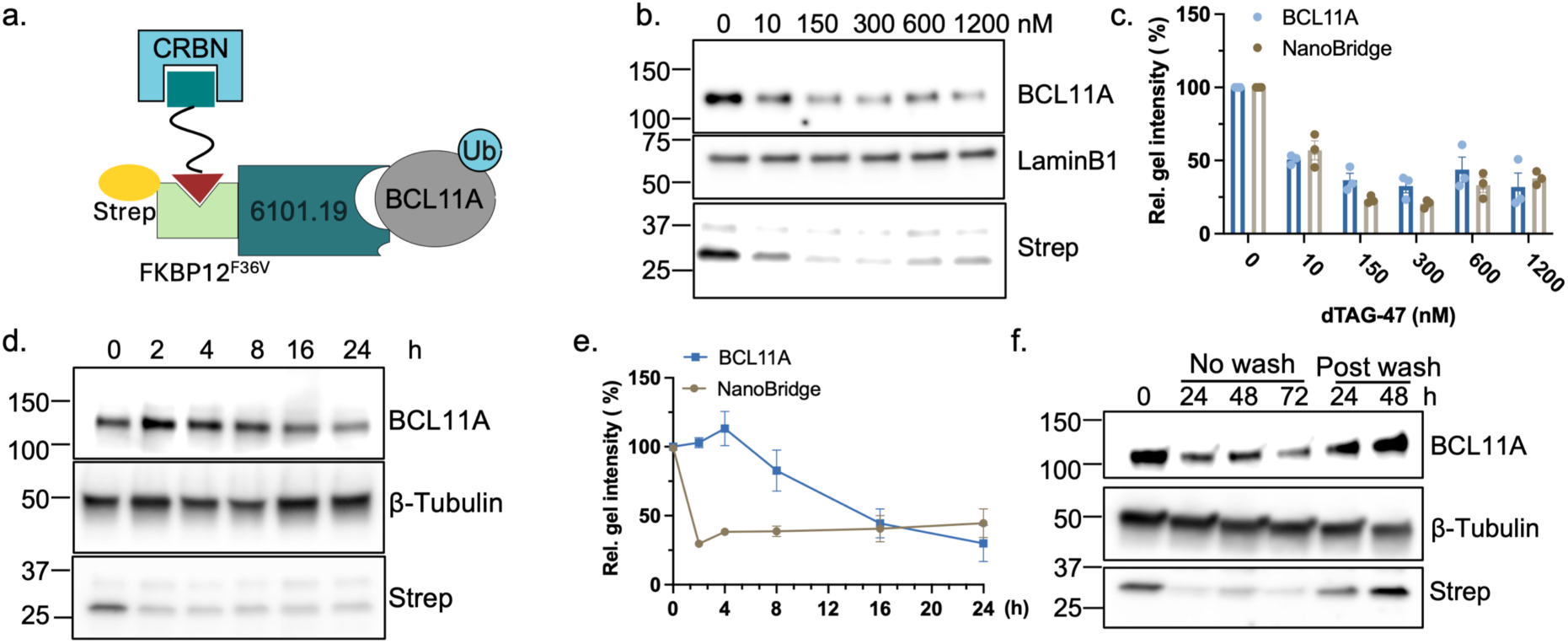
NanoBridge-induced targeted degradation of BCL11A in HUDEP-2 cells. (a) Schematic overview of NanoBridge strategy to induce targeted degradation of BCL11A through the recruitment of CRBN. (b) Immunoblots revealing a dose-dependent BCL11A degradation in cells expressing Nb6101.19-FKBP12F36V. Cells were treated with increasing concentrations of dTAG-47 as indicated for 24 h. (c) Quantification of immunoblots signal of BCL11A relative to Lamin-b1 shown in (b) (mean ± s.d., n = 3). (d) Immunoblots revealing a time-dependent BCL11A degradation in cells expressing Nb6101.19-FKBP12F36V. Cells were treated with 300 nM dTAG-47 for the indicated time. (e) Quantification of immunoblots signal of BCL11A relative to β-tubulin shown in (d) (mean ± s.d., n = 3). (f) Immunoblots revealing removal of dTAG-47 for the indicated time resulted in the recovery of BCL11A. Cells were treated with 300 nM dTAG-47 for 24 h to 72 h. For the washed groups, cells were washed with PBS after 24 h of dTAG-47 treatment and cultured in fresh medium for another 24 or 48 h.

Given that BCL11A is a key repressor of fetal hemoglobin, we further investigated the effect of its degradation on fetal hemoglobin reactivation in primary CD34^+^ progenitor cells isolated from human blood. The CD34^+^ cells were cultured under differentiation conditions and transduced with a lentivirus on Day 3; puromycin was added on Day 5 to select for cells stably expressing Strep-Nb6101.19-FKBP12^F36V^-NLS. These cells and control cells lacking the NanoBridge were then treated with 300 nM dTAG-47 on Day 8 of differentiation (**Figure 3a**). Compared to cells without NanoBridge expression, decreasing levels of BCL11A from Day 9 and reactivation of γ-hemoglobin from Day 10 were marked in cells expressing Nb6101.19-FKBP12^F36V^-NLS (**Figure 3b**). The increased expression of fetal hemoglobin was also observed by qRT-PCR, which revealed a 4.3-fold increase in γ-hemoglobin (HBG) transcript levels only upon dTAG-47 treatment. In contrast, the absence of dTAG-47 resulted in no change to BCL11A levels, even with the NanoBridge present (**Figure S1**).

**Figure 3.**
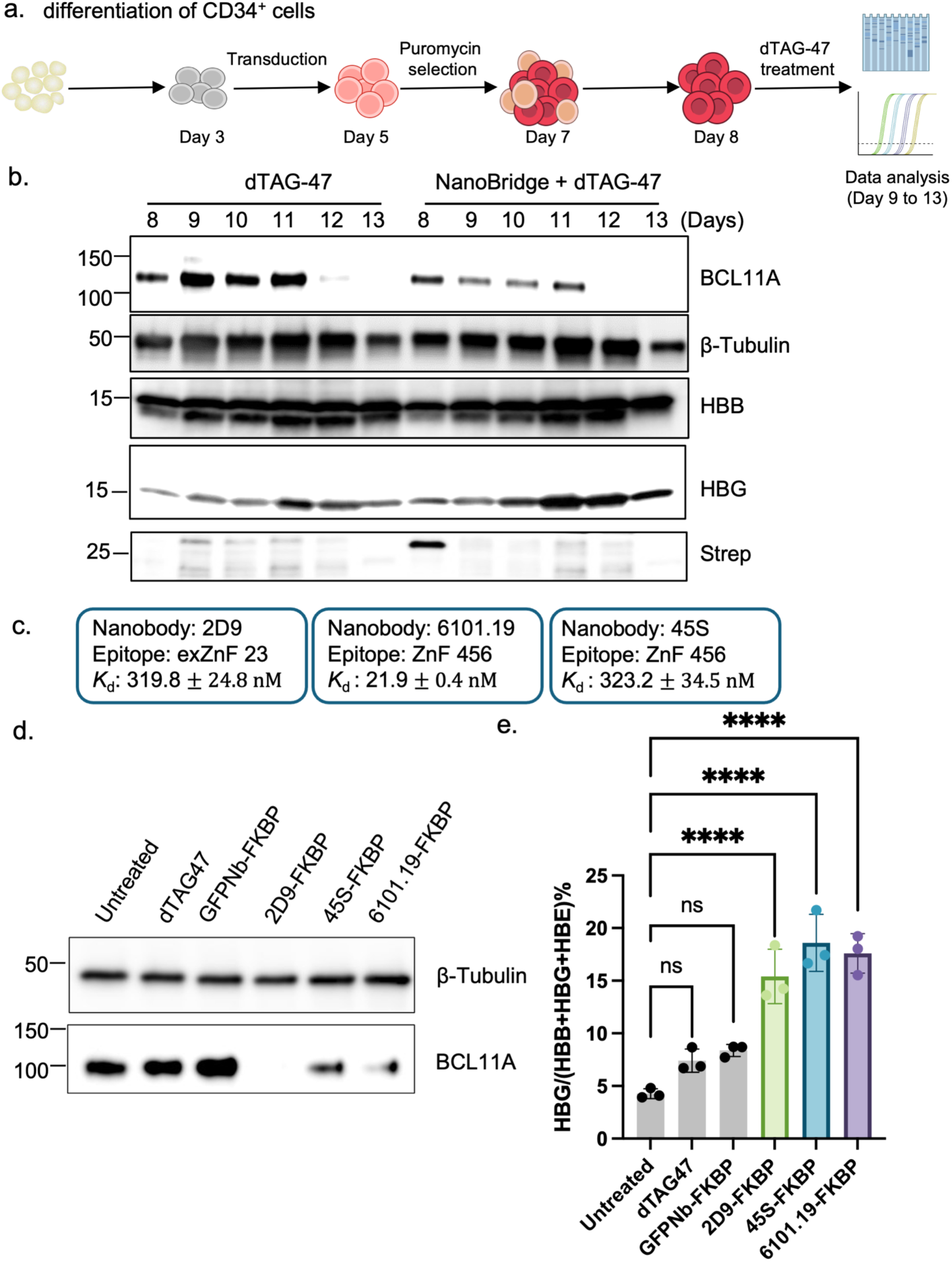
NanoBridge-induced targeted degradation of BCL11A and reactivation of γ-globin in CD34+ cells. (a) Schematic overview of the generation of NanoBridge-expressing CD34+ cells and the treatment of dTAG-47 during cell differentiation. (b) Immunoblots revealing degradation of BCL11A induced by the NanoBridge system during the differentiation of CD34+cells. CD34+ cells with or without Nb6101.19-FKBP12F36V were treated with 300 nM of dTAG-47 on Day 8, and samples from days 9, 10, 11, 12 and 13 were analyzed. (c) Reported binding affinities and epitopes of BCL11A specific nanobodies that were used for NanoBridge constructs. (d) Immunoblots revealing BCL11A level and (e) qRT-PCR showing mRNA levels in CD34+ cells with or without expressing NanoBridge constructs, including GFPNb-FKBP12F36V, Nb2D9-FKBP12F36V, Nb45S-FKBP12F36V and Nb6101.19-FKBP12F36V. CD34+ cells were treated with or without dTAG-47 on Day 8, samples from Day 13 were collected and analyzed (mean ± s.d., n = 3, ****P < 0.0001).

Next, we explored if and how the epitopes and binding affinities of nanobodies influence the degradation efficiency. To do this, we created NanoBridges employing two additional BCL11A nanobodies that were both reported to bind with *K*_d_s approximately 15-fold weaker than Nb6101.19. These two nanobodies bind to different epitopes: nanobody 2D9 (Nb2D9) binds to an extended region surrounding ZnF23 with a *K*_d_ of 319.8 ± 24.8 nM^29^ and nanobody 45S (Nb45S)^37^, a D45S variant of Nb6101.19, binds to ZnF456 with a *K*_d_ of 323.2 ± 34.5 nM. As a negative control, a GFP binding nanobody (GFPNb) with a *K*_d_ of 1.4 nM^38^ was fused to FKBP12^F36V^ to create a NanoBridge that should have no affinity for BCL11A. Cells expressing all three BCL11A-targeting NanoBridge constructs showed significant BCL11A loss upon dTAG-47 treatment when compared to the DMSO-treated group (**Figure 3d**). In contrast, cells without NanoBridge or those expressing the GFPNb-FKBP12^F36V^ did not show such an effect after dTAG-47 treatment, confirming that the degradation is dependent on target-specific engagement. Interestingly, the three BCL11A NanoBridges all showed similar degradation efficiency, suggesting that there is little correlation between binding affinity and degradation efficiency. Indeed, Nb2D9 that binds to extended mediated the highest levels of BCL11A degradation (near depletion) despite having the weakest binding to the target. Consequently, all three BCL11A-binding NanoBridges induced comparable increases in the mRNA levels of HBG upon dTAG-47 treatment; no significant changes were detected in cells lacking the BCL11A NanoBridge or in cells expressing GFPNb-FKBP12^F36V^ (**Figure 3e**). Taken together, these data provide strong support that the degradation of BCL11A and reactivation of HBG induced by the NanoBridge system is dependent on binding of BCL11A nanobodies, and that the platform is accommodating of different epitopes and POI binding affinities.

### NanoBridge induced targeted degradation of KRAS

Encouraged by the successful degradation of BCL11A, we next explored the generalizability of NanoBridge-mediated degradation for other proteins. Kirsten rat sarcoma viral oncogene homologue (KRAS) mutations are present in approximately 25% of all tumors, acting as a primary “driver” of many cancers, such as non-small cell lung cancer, colorectal cancer, and pancreatic cancer.^39^ KRAS has long been deemed a challenging therapeutic target due to its smooth spherical structure lacking deep surface pockets for small molecules; only very recently have small-molecule KRAS inhibitors been reported.^40^ However, Designed Ankyrin Repeat Proteins (DARPins) K27^41^ and K19^42^ bind to KRAS with high affinity and variable specificity. We reasoned that these could be used in the NanoBridge platform to degrade KRAS.

DARPin K27 is a pan RAS binder that engages with the inactive RAS-GDP form; it displays a *K*_d_ of 3.9 nM for its epitope. Meanwhile, DARPin K19 specifically binds to the KRAS isoform with a *K*_d_ of 9.9 nM. To determine if KRAS proteins can be targeted for proteasomal degradation by the NanoBridge system, we generated four NanoBridge constructs where K27 or K19 were fused to the N and C-termini of FBP12^F36V^ (**Figure 4a**) and transduced into HEK293T cell lines to stably express these proteins. Following this, the cells were treated with dTAGs, and KRAS degradation was observed only in cells expressing the RAS-targeting NanoBridges. In these experiments, the VHL-recruiting dTAG^V^-1 appeared to induce KRAS degradation to a greater degree than the CRBN-recruiting dTAG-47 (**Figure 4b**). Interestingly, studies using conventional binder-enzyme direct fusion, where DARPins were fused to an E3 ligase, reported degradation only with VHL fused to the N-terminus of the DARPins, suggesting that direction fusions are sensitive to different orientations of the degrader.^43^ Our data shows that the NanoBridge system instead has no such orientation preference.

**Figure 4.**
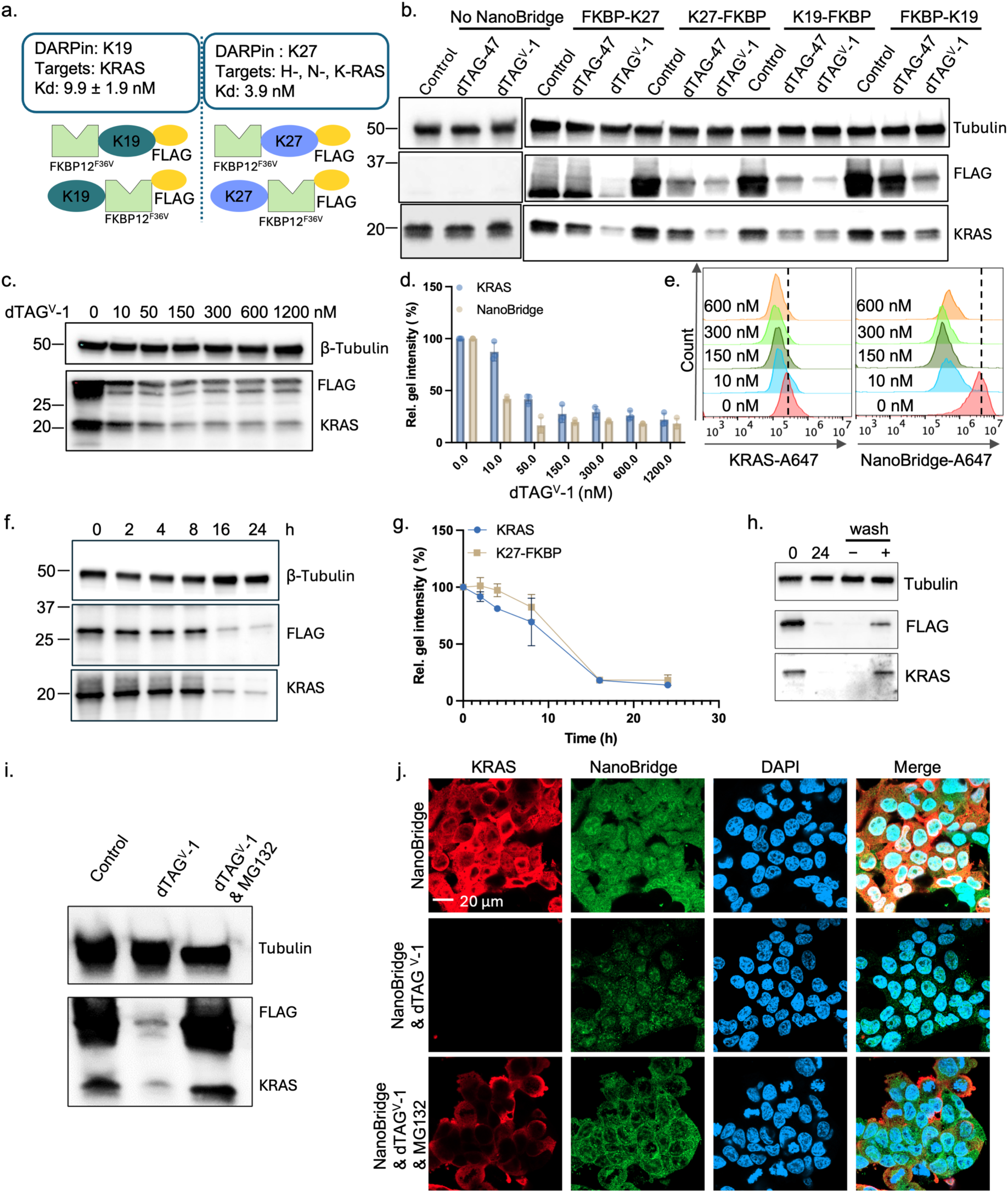
NanoBridge-induced targeted degradation of KRAS in HEK293T cells. (a) Schematic overview of NanoBridge constructs adopting DARPins K27 and K19 for targeted degradation of KRAS. (b) Immunoblots revealing degradation of KRAS induced by different NanoBridge constructs. Cells with or without NanoBridge constructs were treated with 1μM dTAG-47 or dTAGV-1 for 24 h. (c) Immunoblots revealing a dosedependent KRAS degradation in cells expressing K27-FKBP12F36V. Cells were treated with increasing concentrations of dTAGV-1 as indicated for 24 h. (d) Quantitation of immunoblot signal of KRAS relative to β-tubulin shown in (c) (mean ± s.d., n = 3). (e) Flow cytometric analysis of immunostained cells showing a dose-dependent KRAS degradation in cells expressing K27-FKBP12F36V. Cells were treated with increasing concentrations of dTAGV-1 as indicated for 24 h. (f) Immunoblots revealing a time-dependent KRAS degradation induced by NanoBridge system in cells expressing K27-FKBP12F36V. Cells were treated with 400 nM dTAGV-1 for the indicated time. (g) Quantification of immunoblot signal of KRAS relative to β-tubulin shown in (f) (mean ± s.d., n = 3). (h) Immunoblots revealing removal of dTAGV-1 resulted in the recovery of KRAS in 24 h. (i) Immunoblots revealing KRAS degradation is dependent on proteosome. (j) Confocal microscopy images showing the degradation of KRAS (red) induced by NanoBridge (green) is dependent on a functional proteosome. Blue indicates nucleus. Scale bar = 20 μm.

With K27-FKBP12^F36V^ expressing cells, we further evaluated the dose response and kinetics of KRAS degradation. Similarly to BCL11A, treatment with dTAG^V^-1 led to a dose-dependent degradation of KRAS in HEK293T cells, with maximum degradation achieved at concentrations greater than 150 nM (**Figures 4c-e**). Additional studies showed that KRAS was degraded 4 h after treatment with dTAG^V^-1, with the maximum degradation observed after 16 h. Interestingly, unlike what was observed with BCL11A, the rates of K27-FKBP12^F36V^ and KRAS degradation are similar. This suggests that the intrinsic properties of the POI and its interaction with the NanoBridge and E3 ligases impact the degradation efficiency in ways that are not yet known. KRAS levels remained low for at least 48 h, and removal of dTAG^V^-1 via a washout showed a recovery to basal levels within 24 h (**Figure 4f**). Furthermore, treatment with the proteasome inhibitor MG-132 prevented the degradation of both KRAS and NanoBridge, as observed via immunoblotting (**Figure 4i**) and immunofluorescence confocal microscopy (**Figure 4j**). Together, these data confirmed that the NanoBridge-mediated loss of KRAS occurred via proteasomal degradation.

We also explored the effects of the NanoBridge system in multiple cancer cells expressing different KRAS mutations. We first evaluated the degradation in pancreatic cancer Mia PaCa-2 cells that bear KRAS with a G12C mutation. Mia PaCa-2 cells stably expressing four DARPin-FKBP12^F36V^ constructs were obtained and treated with 1 μM dTAG-47 or dTAG^V^-1 for 24 h. Immunoblotting revealed that the two K19 NanoBridges induced greater KRAS degradation than the K27 constructs, perhaps due to the higher expression of the K19 constructs in cells (**Figure S2a**). We also noted that both dTAG-47 and dTAG^V^-1 performed similarly in terms of the degradation efficiency, which differs from what was observed with wild-type KRAS in HEK293T cells (**see Figure 4b**). Similar experiments were performed using K19-FKBP12^F36V^ in the colon cancer cell line HCT116 carrying the G13D KRAS variant, and in the lung cancer cell line A549 that expresses KRAS G12S. Results from these experiments revealed the degradation of KRAS variants with both dTAG-47 and dTAG^V^-1 treatment (**Figures S2b, S2c**). Together, the data show that the NanoBridge system is readily expandable for the degradation of different proteins.

### Targeted phosphorylation and acetylation of p53

A key advantage of the NanoBridge is its potential to recruit different endogenous enzymes to install diverse PTMs on a POI by simply switching the heterobifunctional small molecules. The tumor suppressor p53^44^ is an ideal model for this application, as it has been reported to undergo more than 200 PTMs that collectively form a complex “PTM code” governing p53 stability, its ability to bind DNA, and downstream cell fate decisions.^45^ Among these modifications, phosphorylation^46^ and acetylation^47,48^ are thought to play central roles in regulating p53 stress response and tumor suppressive transcriptional activities.

To evaluate the versatility of the NanoBridge for inducing different types of PTMs, we used ALFA-tagged p53 as a model substrate, wherein the ALFA tag (SRLEEELRRRLTE) was fused to the N-terminus of p53; this protein was stably expressed in H1299, a non-small cell lung cancer cell line, where the basal wild-type p53 levels are negligible. With the high-affinity anti-ALFA nanobody (NbAF, *K*_d_ = 26 pM), we generated the NanoBridge construct NbAF-FKBP12^F36V^. The Choudhary group previously reported PHICS that recruit distinct kinases to induce targeted phosphorylation,^18,19,49^ whereas the Parker group developed AceTAG-1, an acetylation-tagging system that acetylates FKBP12^F36V-^POI fusions^14,15^. Treatment of cells expressing the ALFA-p53 and NbAF-FKBP12^F36V^ with 5 μM PHICS3.3 (which recruits AMPK) induced a 1.9-fold increase in phosphorylation of p53 at S15 (**Figures 5a-c**); simply switching the small molecule to AceTAG-1 induced a 3.2-fold increased acetylation at K382 (**Figure 5a, d, e**).^50^ These results demonstrated that a single NanoBridge construct can induce different types of PTMs when supplied with different heterobifunctional ligands.

**Figure 5.**
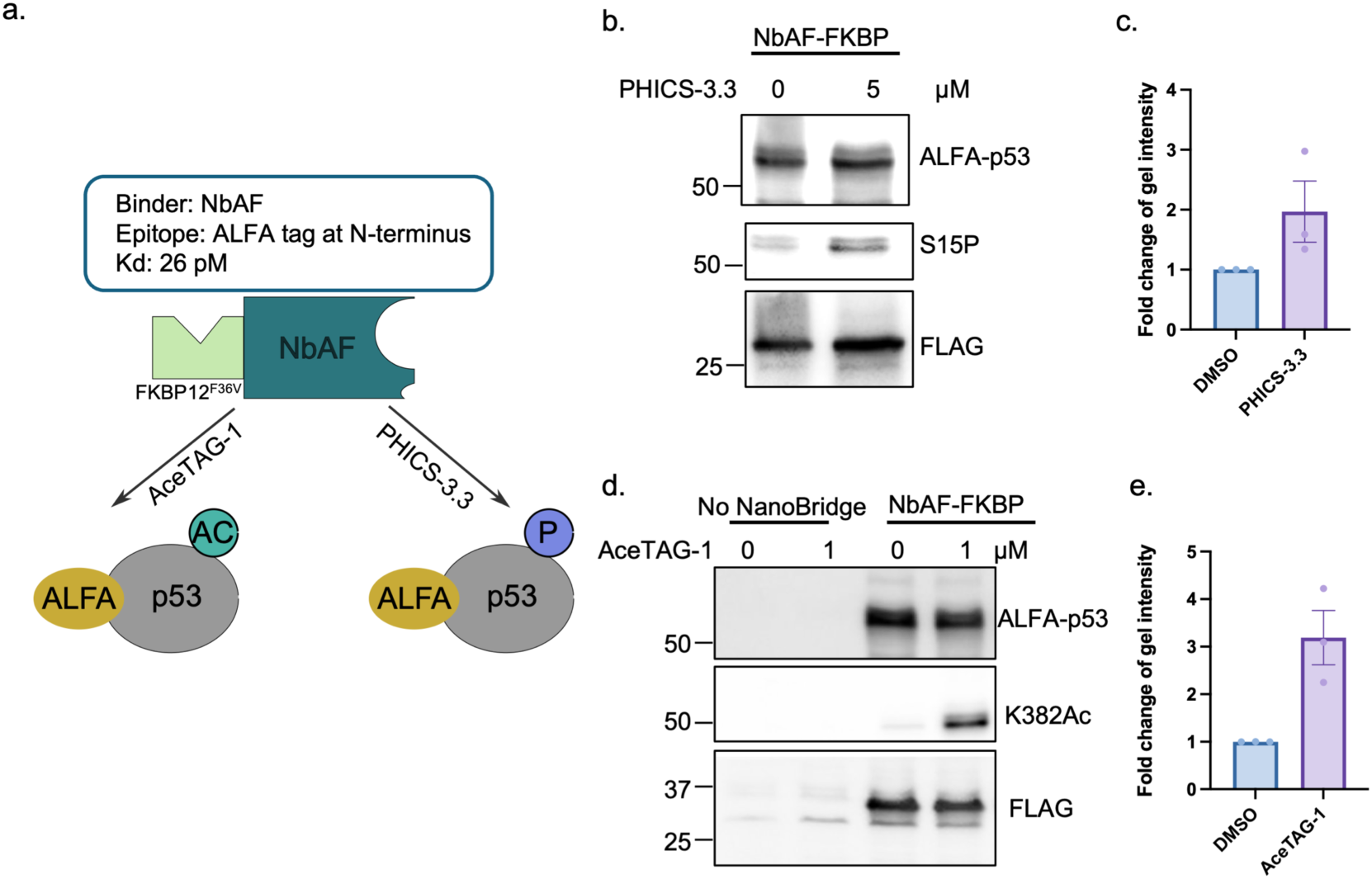
NanoBridge induced phosphorylation and acetylation of ALFA-tagged p53. (a) Schematic overview of NanoBridge construct adopting NbAF for targeted phosphorylation and acetylation of ALFA-p53. Heterobifunctional molecules AceTAG-1 (binding to FKBP12F36V and p300/CBP) and PHICS-3.3 (binding to FKBP12F36V and AMPK) were used. (b) Immunoblots revealing phosphorylation at site S15 of ALFA-p53 in H1299 cells expressing NbAF-FKBP12F36V and ALFA-p53 was treated with PHICS-3.3 at the indicated concentration for 6h. (c) Quantification of immunoblot intensity of phosphorylation relative to FLAG shown in (b) (mean ± s.d., n = 3). (d) Immunoblots revealing acetylation at site K382 of ALFA-p53 in H1299 cells expressing NbAF-FKBP12F36V and ALFA-p53 was treated with AceTAG-1 at the indicated concentration for 16h. (e) Quantitation of immunoblot intensity of acetylation relative to FLAG shown in (d) (mean ± s.d., n = 3).

Encouraged by these results, we further tested if the PTM modification can be achieved on an unmodified/tagless p53. While p53 contains substantial structural disorder, it does have defined domains. There is an N-terminal transactivation domain (TAD, residues 1–61), a proline-rich domain (PRD, residues 64–92), a DNA-binding domain (DBD, residues 96–292) linked to a tetramerization domain (TD, residues 324–356), and a C-terminal regulatory domain (CTD, residues 364–393) (**Figure 6a**). Two protein binders, nanobody 139 (Nb139)^51^ and DARPin C10^52^, have been reported to bind to the structurally ordered DBD of p53 with *K*_d_s of 1 µM and 243 nM, respectively. By fusing FKBP12^F36V^ to these two binders, we generated two NanoBridge constructs (Nb139-FKBP12^F36V^ or C10-FKBP12^F36V^). As before, lentiviral transduction was performed to allow the stable expression of wild-type unmodified p53 and Nb139-FKBP12^F36V^ or C10-FKBP12^F36V^ in H1299 cells. With these cells, AceTAG-1 treatment induced significant K382 acetylation at concentrations as low as 250 nM for both Nb139-FKBP12^F36V^ and C10-FKBP12^F36V^ (**Figure 6b**). A slight “hook effect” was observed for Nb139-FKBP12^F36V^ at AceTAG-1 concentration of 2 μM; this was not observed for the C10-FKBP12^F36V^ NanoBridge.

**Figure 6.**
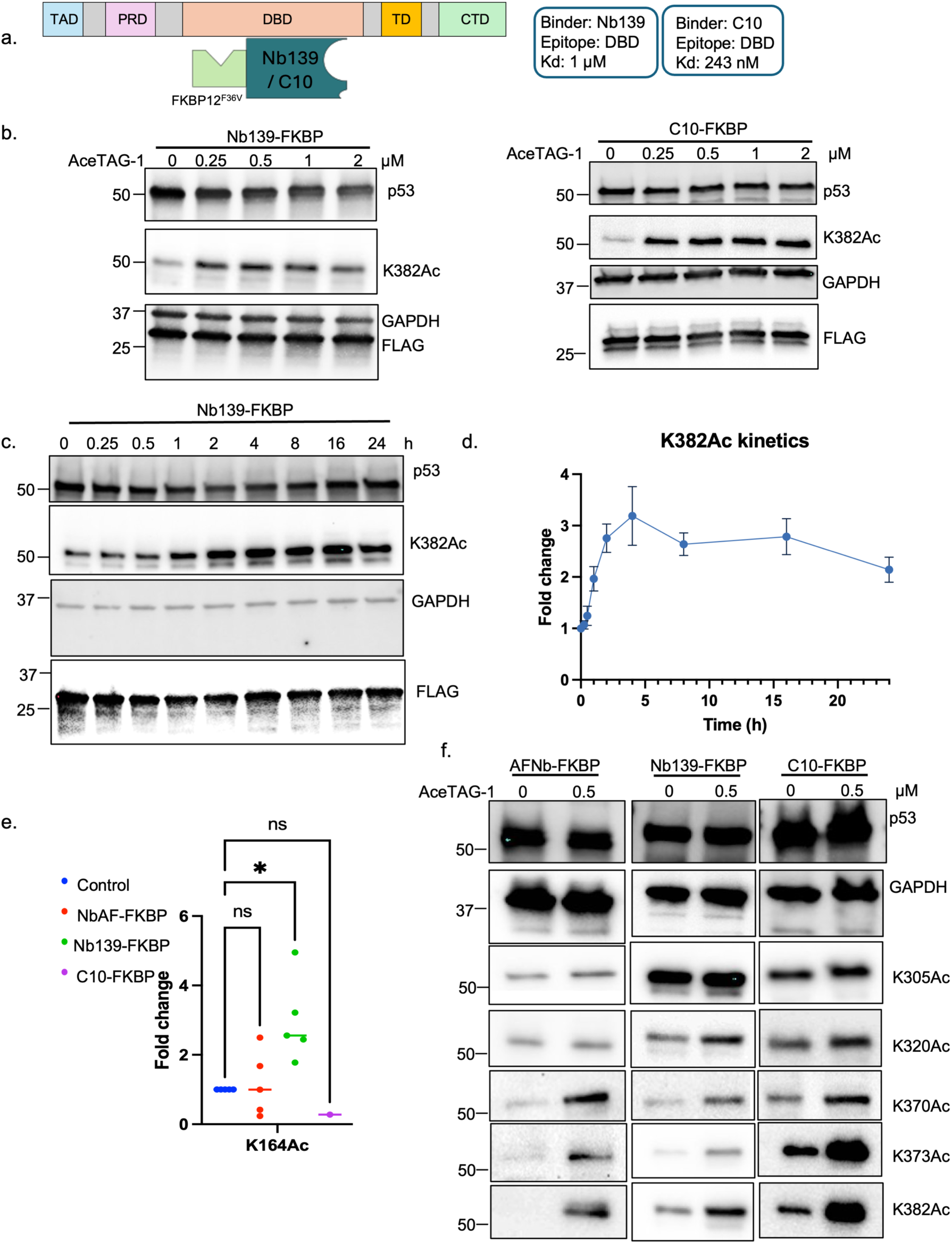
NanoBridge-induced targeted acetylation of p53 in H1299 cells. (a) Schematic overview of NanoBridge constructs using Nb139 and DARPin C10 for targeted acetylation of p53. (b) Immunoblots revealing acetylation of p53 induced by NanoBridge constructs Nb139-FKBP12F36V and C10-FKBP12F36V. Cells expressing NanoBridge constructs were treated with increasing concentrations of AceTAG-1 for 24h. (c) Immunoblots revealing the kinetics of p53 acetylation at site K382. Cells expressing Nb139-FKBP12F36V were treated with 1 μM AceTAG-1 for the indicated times. (d) Quantitation of immunoblot signal of K382Ac relative to GAPDH shown in (c) (mean ± s.d., n = 3). (e) Proteomics analysis revealing acetylation sites on p53 induced by NanoBridge systems. The abundance of acetylated peptide was normalized to total peptide, and fold change to the DMSO treated group was calculated (mean ± s.d., n = 4). Note: acetylated lysines may be detected from different peptide isoforms; each dot represents the fold change from a same peptide isoform. (f) Immunoblots revealing acetylation sites on p53 induced by NanoBridge systems. Cells expressing NbAF-FKBP12 F36V, Nb139-FKBP12F36V and C10-FKBP12F36V were treated with 0.5 μM AceTAG-1 for 16h.

A time course experiment to monitor the kinetics of acetylation at K382 showed that a significant increase in acetylation was detectable after ∼30 min of AceTAG-1 treatment (**Figures 6b, c**), much faster than that determined for degradation (**Figures 2e, 4g**) and potentially influenced by the intrinsic enzyme activity difference between E3 ligases and acetyltransferases. As with the degradation experiments, removal of AceTAG-1 via washout led to the loss of acetylation at this position in 12 h (**Figure S3**).

To more globally map the p53 acetylation sites that are induced with the NanoBridge platform, we systematically compared the acetylation patterns of cells expressing NbAF-FKBP12^F36V^, Nb139-FKBP12^F36V^, and C10-FKBP12^F36V^. Using quantitative mass spectrometry, p53 from cells with and without treatment of AceTAG-1 was immunoprecipitated, digested with trypsin, and the acetylated peptides were quantified via tandem mass spectrometry. These data revealed a substantial increase in acetylation at K164 (a site known to be acetylated by p300/CBP) for construct Nb139-FKBP12^F^^36^; a similar increase in the acetylation at that site was not detected for the other two NanoBridges (NbAF-FKBP12^F36V^ and C10-FKBP12^F36V^, **Figures 6d, S4**). Peptides carrying acetylation at sites K292, K305, K320, and K357 were also detected, but their abundances were statistically similar for all three constructs.

It is known that p53 acetylation at the C-terminal regulatory domain is challenging to detect via proteomics due to the multitude of trypsin cleavage sites in that domain. As such, we performed immunoblotting with antibodies that recognize specific lysine acetylation sites of p53 instead. After treatment with AceTAG-1, all three NanoBridge constructs resulted in a significant acetylation increase at sites K370, K373, and K382, which are all known to be sites modified by p300/CBP (**Figure 6f**). Consistent with the mass spectrometry results (**Figure S4**), no significant changes were observed at sites K305 and K320, although K305 is a known site of p300/CBP. These data confirmed that the NanoBridge induced proximity led to the acetylation on multiple endogenous sites of p300/CBP that include K164, K370, K373, and K382. For K320, the predominant site of acetyltransferase PCAF/GCN5, we did not see such a significant increase. It is therefore possible that the site preference of acetylation induced by NanoBridge is influenced by the inherent biological preferences of the recruited PTM writer. Interestingly, only Nb139-FKBP12^F36V^ induced K164 acetylation, suggesting that difference in binding affinity or epitopes may lead to distinct acetylation patterns and levels.

Collectively, these data established that the NanoBridge system can induce rapid, efficient, and reversible acetylation on multiple sites of an intrinsically disordered protein without the need for genetically encoded fusion tags. We envision that this system can serve as a useful strategy to gain deeper insights into the functional roles of PTMs on endogenous proteins.

## Conclusion and discussion

In this work, we report the NanoBridge platform as a highly modular platform to induce the targeted modulation of proteins via degradation, phosphorylation, and acetylation of traditionally undruggable POIs in live cells. The NanoBridge consists of a protein binder, such as nanobody or DARPins, fused with a FKBP12^F36V^ tag. By leveraging different FKBP12^F36V^ targeting heterobifunctional small molecules, the NanoBridge enables the recruitment of multiple distinct endogenous PTM enzymes to achieve different types of targeted PTMs on POIs in cells. As an example, two E3 ligases recruiting heterobifunctional small molecules (dTAG-47 and dTAG^V^-1) allowed for the targeted degradation of BCL11A and KRAS. Furthermore, by simply changing these dTAG molecules for kinase recruiter (PHICS-3.3) and acetyltransferase recruiter (AceTAG-1), we demonstrated targeted phosphorylation and acetylation on p53. With AceTAG-1, we further demonstrated that the targeted acetylation can be achieved on p53 without requiring genetic fusion tags. Our results indicate that the NanoBridge system is highly modular, flexible, and enables multiple types of targeted PTMs on difficult-to-drug POIs. Additionally, the use of heterobifunctional small molecules makes NanoBridge-induced PTMs reversible and allows for precise temporal control in the induction of PTMs.

The ability to edit endogenous proteins in situ is essential for advancing our understanding of their roles and for developing therapeutics. The CIP paradigm has transformed this field by enabling specific protein modulation without perturbing enzyme activity or affecting global substrates. However, a large fraction of the human proteome remains challenging to target with small-molecule ligands. Consequently, exogenous tags ranging from relatively small peptides to larger fusion proteins (for example, GFP^53^) are often used to provide a handle on the target POIs. While these tags have greatly advanced the study of proteins in cells, they can influence the intrinsic properties of target proteins. The NanoBridge platform developed here filled this gap by allowing the direct editing of tag-free POI in cells. Unlike the traditional enzyme-protein direction fusions, the NanoBridge provides more flexibility to induce different types of PTMs by recruiting endogenous PTM enzymes instead of introducing exogenous versions that could perturb original PTM homeostasis and may generate nonphysiological modifications. As exemplified by the p53 acetylation using p300/CBP recruiting AceTAG-1, there is a rapid and efficient acetylation at multiple sites of p53 using the NanoBridge. And all the modification sites we observed (K164, K370, K373, and K382) are known endogenous sites of p300/CBP. Interestingly, of the three nanobridges studied, only Nb139-FKBP12^F36V^ induced K164 acetylation, indicating that binding affinity and epitopes of biologic binder may lead to correspondingly distinct acetylation patterns. As protein binders have the flexibility to target different epitopes on the POI, we envision that the NanoBridge platform may serve as a versatile tool to investigate functional consequences of acetylation in cells, particularly when comparing distinct acetylation patterns to gain deeper insights into their functions.

Despite these advantages, the NanoBridge system has several limitations. First, the NanoBridge constructs have only been expressed in cells. This limitation can be addressed with the advancement of protein delivery methods, including those we have evaluated in prior studies.^54–57^ Second, the binding of a nanobody or DARPin may influence the function of POIs. These include preventing or restricting protein–protein interactions, inhibiting enzymatic activity, or interfering with PTM dynamics. We posit that additional validation experiments with appropriate controls (for example, using the nanobody alone to only induce binding) may reveal the degree to which these functions are impacted. Third, the application range of the NanoBridge system is determined by the availability and efficiency of proximity-inducing small molecules. We only demonstrated ubiquitination, phosphorylation, and acetylation here but predict that future studies will enable use of the NanoBridge for the installation or removal of additional types of PTM.

In conclusion, we demonstrated the NanoBridge as a generalizable and flexible strategy to mediate the targeted degradation, phosphorylation, and acetylation of proteins in live cells. This strategy enables the FKBP12^F36V^ binding small molecules to transiently induce the proximity of endogenous PTM enzymes to target proteins, thus facilitating the PTM editing on proteins in their native forms. NanoBridge-induced PTM editing is target-specific, efficient, reversible, and has temporal control. Moreover, it enables switching between different PTM types and supports multisite PTM editing, capturing the dynamic nature of PTM regulation. It therefore represents a useful platform for the targeted modulation of endogenous proteins and provides new opportunities to investigate protein PTMs for biological discovery.

## Acknowledgements

This work was supported in part by NIH grant R01HL173127 (to L.M.K.D.). L.M.K.D. was supported by a Stanford Terman Fellowship and a MAC3 Impact Philanthropies Faculty Fellowship at the Sarafan ChEM-H Institute. We thank Dr. Christopher Parker for providing AceTAG-1 compound and advice on p53 proteomics. We are grateful to Dr. Amit Choudhary for providing the PHICS molecule and for general advice on p53 phosphorylation. The generation of PHICS-3.3 was supported by NIH grant R01DK132900 (to A.C.). We acknowledge the Stanford University Cell Sciences Imaging Core Facility (RRID: SCR_017787) for confocal microscope access. We thank Dr. Gabriela Grigorean for performing the proteomics in the Proteomics Core Facility of the Genome Center at the University of California, Davis. The Bruker timsTOF HT LC/MS system was supported by the Howard Hughes Medical Institute, Investigator Award for Dr. Neal Hunter, UC Davis. We thank Profs. Peter Kim and Jonathan Long for access to equipment.

## Note

The authors declare no competing financial interest.

## 1. Materials and Methods

### Cell culture

Cells were cultured in a humidified atmosphere of 5% CO_2_ and 95% air at 37°C. Mycoplasma tests were carried out routinely to detect contamination. For cell counting and viability measurement, the cell suspension was mixed 1:1 with 0.4% trypan blue dye (Thermo Scientific, #15250061) and measured with Countess II FL Automated Cell Counter (Thermo Scientific).

HUDEP-2 cells (RCB4557) were obtained from Riken Bio Resource Research Center (Japan). Cells were cultured according to the reported method.^1^ In brief, cells were maintained in expansion medium, which contains StemSpan serum-free expansion medium (SFEM, StemCell Technologies, #09650), 2% Penicillin-Streptomycin solution (10,000 U/mL stock), 50 ng/mL recombinant human stem cell factor (SCF, StemCell Technologies, #78062.2), 3 IU/mL Epoetin alfa (EPO, Sigma, #H5166), 0.4 μg/mL dexamethasone (Sigma, #D4902), and 1 μg/mL doxycycline (Thermo Scientific, # BP26531).

Peripheral blood-derived CD34^+^ cells from multiple donors were obtained from the NIDDK-Center of Excellence in Hematology at the Fred Hutchinson Cancer Research Center. The cells were cultured according to the reported method.^2^ In brief, cells were cultured in erythroid differentiation medium (EDM), which contains IMDM medium (Gibco, #12440053) supplemented with stabilized glutamine, 330 ug/mL holo-human transferrin (Sigma, #T0665), 10 ug/mL recombinant human insulin (Sigma, #I9278), 2 IU/mL heparin (StemCell Technologies, #07980), and 5% inactivated human plasma type AB (Sigma, #H5667). The expansion procedure comprised 3 steps. In the first step (Day 0 to Day 7), 10^4^/mL CD34^+^ cells were cultured in EDM in the presence of 10^-6^ M hydrocortisone (StemCell Technologies, #07925), 100 ng/mL SCF, 5 ng/mL IL-3 (StemCell Technologies, #78146), and 3 IU/mL EPO. On Day 4, 1 volume of cell culture was diluted in 4 volumes of fresh medium containing SCF, IL-3, EPO, and hydrocortisone. In the second step (Day 7 to Day 11), the cells were resuspended at 10^5^/mL in EDM supplemented with SCF and EPO. In the third step (Day 11 to Day 13), the cells were cultured in EDM supplemented with EPO alone. Cell counts were adjusted to a range of 7.5 × 10^5^ - 1 × 10^6^ on Day 11.

Human HEK293T cells (female) were purchased from ATCC. The cells were cultured in DMEM supplemented with 10% FBS, 1% Penicillin-Streptomycin solution (10,000 U/mL stock).

MIA PaCa-2 cells (CRL-1420) were purchased from ATCC. The cells were cultured in DMEM supplemented with 10% FBS, 2.5% horse serum, and 1% Penicillin-Streptomycin solution (10,000 U/mL stock).

HCT116 cells (CCL-247) were purchased from ATCC. The cells were cultured in McCoy’s 5a Medium Modified (Thermo Scientific, #16600082), supplemented with 10% FBS, and 1% Penicillin-Streptomycin solution (10,000 U/mL stock).

A549 cells was a gift from Dr. Jiang Hong Rao lab at Stanford University School of Medicine. The cells were cultured in DMEM, high glucose (Thermo Fisher Scientific, #11965), supplemented with 10% FBS and 1% Penicillin-Streptomycin solution (10,000 U/mL stock).

H1299 cells was a gift from Dr. Maximillian Diehn lab at Stanford University School of Medicine. The cells were cultured in RPMI 1640 Medium (Thermo Scientific, #11875093), supplemented with 10% FBS and 1% Penicillin-Streptomycin solution (10,000 U/mL stock).

### Plasmid construction

All gene blocks were obtained from GENEWIZ while primers were obtained from Integrated DNA Technologies. KOD hot start polymerase (Sigma, #71086-3) was used for PCR reaction, the primers used are listed in Table S1. The PCR products was digested with DPNI (NEB, #R0176L) and purified with PCR purification kit (Invitrogen, #K310001). NEBuilder HiFi DNA Assembly (NEB, #E2621X) was used for plasmid fusion.

### Transformation

2 µL of plasmid (concentration ≥20 ng/µL) was added to a thawed tube of Stellar competent cells (Takara, #636763) and incubated on ice for 15 min. The cells were heat shocked at 42 ℃ for 45 s and placed on ice for 2 min. 900 µL of LB media were added and the cells cultured for 1 h at 37 ℃ while shaking at 200 rpm. Cells were centrifuged for 3 min at 2500 × g and 900 µL LB media was removed. The recovered cells were resuspended in 100 µL remaining media and added to ampicillin-(100 mg/mL) or kanamycin-containing (50 mg/mL) agar plates and incubated at 37 ℃ overnight.

### Lentivirus productions and transduction

To generate NanoBridge expressing cell lines, the corresponding NanoBridge sequence were subcloned into the lentiviral vector pLVX-EF1a-IRES-Puro (Addgene, #85132). Lentiviruses were packaged in HEK293T cells as described previously.^3^ In brief, 1.25 µg plasmid with gene of interest, 0.8 µg psPAX2 (Addgene, #12260) and 0.45 µg pMD2.G (Addgene, #12259) were added into 50 µl of Opti-MEM medium (Thermo Scientific, #31985062). At the same time, 7.25 µg PEI (Polysciences Inc) was added into another 50 µl of Opti-MEM medium. After incubation for 3 min, the plasmid containing medium was added into PEI medium, mixed well and incubated for 15 to 25 min. The mixture was added dropwise into a HEK293T cells cultured in 6-well plate. 12 h later after transfection, medium was replaced with fresh complete DMEM medium. After another 48h, the supernatant was collected and filtered through 0.45 µM filter to give the lentivirus. The lentivirus can be used fresh or save in -80°C for later use.

For transduction, 1 mL of lentivirus containing medium was added to cells cultured in 1 mL of complete medium, incubate for 24 - 48 h. After that medium was changed to fresh complete medium, and cells were cultured for another 2 days. After that, puromycin at the concentration of 1 or 2 µM was used to select the stable expressing cells.

For the transduction of CD34^+^ cells, cells were cultured as the method described above. On Day 3, CD34^+^ cells were incubated with lentivirus containing medium at the ratio of 1:1 and incubated for 48h. On Day 4, the medium was changed to complete CD34^+^ differentiation medium. On Day 5, puromycin at the concentration of 1µg/mL was added to select for the NanoBridge stable expressing cells. The treatment of puromycin was continued until the end of this experiment. From Day 8 to Day 13, cells were treated with dTAG47 at the concentration of 500 nM. Cells from Day 8 to Day 13 were harvested and analyzed.

### Immunoblotting

For degradation studies, cells were washed with cold DPBS buffer and lysed using RIPA buffer (VWR, #97063-270) containing 0.1 U Benzonase nuclease (Santa Cruz biotechnology, #sc-391121B) and EDTA-free protease inhibitor cocktail (Bimake, #B14002). For acetylation studies, cells were washed with cold DPBS buffer and lysed using RIPA buffer containing 0.1 U Benzonase nuclease, EDTA-free protease inhibitor cocktail, 5 mM sodium butyric acid, and 10 mM nicotinamide. Cellular debris was cleared by centrifugation at 16,000 g for 20 min at 4°C. Protein extracts were quantified by Pierce BCA Protein Assay Kit (Thermo Scientific, #PI23235) with a Nanodrop One^C^ (Thermo Scientific) according to the manufacturer’s instruction. Protein lysates (∼ 30 µg/lane) were resolved by SDS-PAGE and transferred onto poly vinylidene difluoride (PVDF) membranes. Membranes were blocked with 5% non-fat milk in 0.1% Tween 20/PBS for 1 h. The blots were then probed with the relevant primary antibodies in blocking solution at 4°C overnight with gentle agitation. Membranes were washed 5 min with 0.1% Tween 20 in PBS three times and were incubated with secondary antibodies for 1 h at room temperature. For HRP-conjugated secondary antibody, antigens were detected by addition of Clarity™ Western ECL Substrate (Bio-Rad, #1705060). The membranes were visualized by chemiluminescence or fluorescence on a Bio-Rad ChemiDoc MP Imaging System. Primary antibodies used: BCL11A (Cell Signaling, #75432), StrepMAB-Classic HRP (IBA, #2-1509-001), GAPDH (Santa Cruz Biotechnology, #sc-365062), Lamin b1 (Proteintech, #66095-1), β-tubulin (Proteintech, #66240-1), β-hemoglobin (Sigma Aldrich, #WH0003043M1), g-hemoglobin (Proteintech, #25728-1-AP). Flag tag (Cell signaling, #14793), KRAS (Proteintech, #12063-1-AP), p53 (Santa Cruz Biotechnology, #sc-126), p53-K382Ac (Cell Signaling, #2525), p53-K305Ac (Abcam, #Ab109396), p53-K320Ac (Sigma, #06-1283), p53-K370Ac (Abcam, #Ab183544), p53-K373Ac (Abcam, #Ab62376), p53-K386Ac (Abcam, #Ab52172). Secondary antibodies used: Anti-Mouse-HRP (Abcam, #ab6728), Anti-Rabbit-HRP (Promega, #W4018), IRDye 680RD Goat anti-Mouse IgG Secondary Antibody (LI-COR, #926-68070), IRDye 880RD Goat anti-Rabbit IgG Secondary Antibody (LI-COR, #926-32213).

### RT-qPCR

Quantitative real-time PCR was performed to quantify RNA abundance. For each sample, total RNA was isolated by using the RNeasy Plus Mini Kit (QIAGEN Cat# 74134), followed by cDNA synthesis using the iScript cDNA Synthesis Kit (BioRad, Cat# 1708890). qRT-PCR primers (Table S2) were ordered from Integrated DNA Technologies. Quantitative real-time PCR was performed using the PrimePCR assay with the iTaq universal SYBR Green supermix (BioRad, #1725120) and run on a Biorad CFX384 real-time system (C1000 Touch Thermal Cycler) according to the manufacturer’s instructions. C_q_ values were used to quantify RNA abundance. The relative abundance of the hemoglobin was normalized to a GAPDH internal control by using this equation:

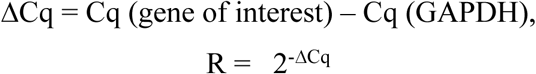

Representative data and calculations are shown in Table S3.

### Confocal imaging

200 µL of HEK293T cells expressing K27-FKBP12^F36V^ were seeded into 8-well chamber slides (Ibidi, #80807) at a density of 2 × 10^5^ cells/mL. Cells were cultured for 24 h at 37 ℃. After that, cells were treated with DMSO, dTAG^V^-1 500 nM or dTAG^V^-1 (500 nM) + MG132 (500 nM) for another 24 h. After treatment, cells were washed with cold PBS buffer once and fixed by addition of 4% paraformaldehyde for 20 min at 4 °C. Then, cells were permeabilized by adding 0.1% Triton-X100 in PBS for 5 min. After that, cells were blocked with 5% BSA in 0.1% Tween 20/PBS for 1h and incubated with anti-FLAG and anti-KRAS antibodies in 1% BSA in 0.1% Tween 20/PBS overnight. Following that, cells were washed with 0.1% Tween 20/PBS for 3 times, and incubated with 1 μg/mL DAPI, secondary antibodies Goat anti-rabbit IgG (H+L) -488 (Invitrogen, #A32731) and Goat anti-Mouse-IgG (H+L) -Alexa Fluor 647 (Invitrogen, #A21235) for 1h at room temperature. Then, cells were washed 3 times with PBST and imaged with a Zeiss LSM980 Confocal Microscope with 60 × oil immersion objective at excitations of 653, 493 and 353 nm.

### Flow cytometry to analyze KRAS degradation

2mL of HEK293T cells expressing K27-FKBP12^F36V^ were seeded into 6-well plate at the density of 2 × 10^5^ cells/mL. Cells were cultured for 24 h at 37 ℃. After that, cells were treated with dTAG^V^-1 at the indicated concentrations for 24 h. After treatment, cells were harvested and washed twice with cold PBS buffer and centrifuged at 350-500 × g for 5 min. Then, cells were fixed with 4% paraformaldehyde for 20 min at 4 °C and permeabilized with 0.2 % Triton-X100 in PBS for 5 min. Following that, 100 μL of 0.1% BSA in 0.1% Tween 20/PBS buffer was added for blocking, and cells were incubated with anti-KRAS or anti-FLAG for 2 h, washed 3 times with 0.1% Tween 20 in PBS buffer and incubated with Goat anti-rabbit -Alexa Fluor 647 (Invitrogen, #A21244) for 1 h at room temperature. After washing 3 times with 0.1% Tween 20/PBS buffer, cells were analyzed on a BD Accuri^TM^ C6 Plus cell sorter and analyzer. The data was analyzed by FlowJo software.

### Immunoprecipitation and proteomics analysis

H1299 Cells were cultured in 10 cm dishes in indicated media. DMSO or AceTAG-1 (500 nM) were added once the cells grew to 70-80% confluency and incubated for 16 h. The cells were harvested and washed with cold PBS twice. The pelleted cells were lysed in 1 mL RIPA lysis buffer, containing 5 mM sodium butyrate, 20 mM nicotinamide, and 1× protease inhibitor (Selleckchem, #B14001), and 0.1 U Benzonase nuclease for 25 min at 4°C. Following that, the cell lysate was centrifugation at 16,000 × *g* for 20 min at 4°C. The protein concentration was normalized after assessing the total protein concentration using the BCA Protein Assay kit (Thermo Scientific, #23227), equivalent amount of protein was subjected to enrichment by p53 antibody. Enrichment was carried out at 4°C for 8h while rotating. During incubation, the magnetic protein A/G beads (Thermo Scientific, #88802) were prewashed with lysis buffer for 3 times. After the primary antibody incubation, magnetic protein A/G beads were added into cell lysate and incubate for another 4h. The beads were enriched by magnetic rack and washed with lysis buffer and cold PBS.

Protein samples on magnetic beads were washed four times with 200 µL of 50 mM Triethyl ammonium bicarbonate (TEAB) with a twenty-minute shake time at 4℃ in between each wash. Roughly 2.5 µg of trypsin was added to the bead and TEAB mixture, and the samples were digested over night at 800 rpm shake speed, at 37°C, followed by a boost addition of trypsin using same wt/wt ratios for overnight digestion at 37°C. To stop the digestion, the reaction mixture was acidified with 1% trifluoroacetic acid (TFA). The supernatant containing the tryptic peptides, was dried in a vacuum centrifuge and re-constituted in water with 2% acetonitrile (ACN).

500 ng total peptide was loaded onto a disposable Evotip C18 trap column (Evosep Biosytems, Denmark) and subjected to nanoLC on a Evosep One instrument (Evosep Biosystems). Tips were eluted directly onto a PepSep analytical column, dimensions: 150 µm x 25 cm C18 column (PepSep, Denmark) with 1.5 μm particle size (100 Å pores) (Bruker Daltronics). Mobile phases A and B were water with 0.1% formic acid (v/v) and 80/20/0.1% ACN/water/formic acid (v/v/v), respectively. The standard pre-set method of 60 samples-per-day (21minute run) was used. The mass spectrometry was done on a hybrid trapped ion mobility spectrometry-quadrupole time of flight mass spectrometer (timsTOF HT, Bruker Daltonics, Bremen, Germany), operated in PASEF mode. The acquisition scheme used for DIA consisted of 36 precursor windows at width of 25m/z, over the mass range 300-1200 Dalton. The TIMS scans layer the doubly and triply charged peptides.

LCMS files were processed with Spectronaut v20 (Biognosys, Zurich, Switzerland) using DirectDIA analysis mode. Mass tolerance/accuracy for precursor and fragment identification was set to default settings. The reviewed FASTA for *Homo sapiens*, UP00005640, downloaded from Uniprot (on 28/02/2025) and a database of 380 common laboratory contaminant proteins (https://www.thegpm.org/crap/) were used. Methionine oxidation and acetylation of protein N termini were set as variable modifications. A decoy false discovery rate (FDR) at less than 1% for peptide spectrum matches and protein group identifications was used for spectra filtering (Spectronaut default). Decoy database hits, proteins identified as potential contaminants, and proteins identified exclusively by one site modification were excluded from further analysis.

For data analysis, the acetylated peptide was normalized to total peptide abundance, and the fold change of normalized acetylated peptide in AceTAG-1 treated group *vs* DMSO treated group was calculated and presented in figure. Note that peptides may not have been detected in every replicate but were required to be detected and quantified in at least two biological replicates to be included in this analysis. And for a same site, different peptide isoforms may be detected and are calculated separately.

### Statistics

All data presented were mean ± SD. Statistical analysis was performed by using Prism 10 and p values obtained from one-way ANOVA, two-way ANOVA or unpaired Student’s t-test where necessary; ns, not significant; *p < 0.05; **p < 0.01; ***p < 0.001; ****p < 0.0001.

## 2. Protein sequences of NanoBridge constructs

### Nb6101.16-FKBP12^F36V^-NLS-Strep tag

MRVQLVESGGGLVQAGGSLRLSCAADGFDFKSYAMGWYRQAPGREDELVAAITASGDYTYYADSVKGRFTISRDNAKNTVYLQMNSLKPDDTAVYYCAALSYVAEGYWGQGTQVTVSSGSGSGSMGVQVETISPGDGRTFPKRGQTCVVHYTGMLEDGKKVDSSRDRNKPFKFMLGKQEVIRGWEEGVAQMSVGQRAKLTISPDYAYGATGHPGIIPPHATLVFDVELLKLEGSKRPAATKKAGQAKKKKGGGSWSHPQFEKGPV

### 2D9-FKBP12^F36V^-NLS-Strep tag

MQVQLVESGGGLVQAGGSLRLSCAASGSIFVNNAMGWYRQAPGKERELVAAISASGGSTYYADSVKGRFTISRDNAKNTVYLQMNSLKPEDTAVYYCAADQDGYPYEYWGQGTQVTVSSGSGSGSMGVQVETISPGDGRTFPKRGQTCVVHYTGMLEDGKKVDSSRDRNKPFKFMLGKQEVIRGWEEGVAQMSVGQRAKLTISPDYAYGATGHPGIIPPHATLVFDVELLKLEGSKRPAATKKAGQAKKKKGGGSWSHPQFEKGPV

### 45S-FKBP12^F36V^-NLS-Strep tag

MRVQLVESGGGLVQAGGSLRLSCAASGFIFDSYAMGWYRQAPGKESELVAAITSSGSSTYYADSVKGRFTISRDNAKNTVYLQMNSLKPEDTAVYYCAALDYVIDGYWGQGTQVTVSSGSGSGSMGVQVETISPGDGRTFPKRGQTCVVHYTGMLEDGKKVDSSRDRNKPFKFMLGKQEVIRGWEEGVAQMSVGQRAKLTISPDYAYGATGHPGIIPPHATLVFDVELLKLEGSKRPAATKKAGQAKKKKGGGSWSHPQFEKGPV

### K19-FKBP12^F36V^-3×FLAG tag

MDLGKKLLEAARAGQDDEVRILMANGADVNASDRWGWTPLHLAAWWGHLEIVEVLLKRGADVSAADLHGQSPLHLAAMVGHLEIVEVLLKYGADVNAKDTMGATPLHLAARSGHLEIVEELLKNGADMNAQDKFGKTTFDISTDNGNEDLAEILQKLGSGSGSMGVQVETISPGDGRTFPKRGQTCVVHYTGMLEDGKKVDSSRDRNKPFKFMLGKQEVIRGWEEGVAQMSVGQRAKLTISPDYAYGATGHPGIIPPHATLVFDVELLKLEASDYKDDDDKGDYKDDDDKGDYKDDDDK

### FKBP12^F36V^-K19-3×FLAG tag

MGVQVETISPGDGRTFPKRGQTCVVHYTGMLEDGKKVDSSRDRNKPFKFMLGKQEVIRGWEEGVAQMSVGQRAKLTISPDYAYGATGHPGIIPPHATLVFDVELLKLEGSGSGSGSDLGKKLLEAARAGQDDEVRILMANGADVNASDRWGWTPLHLAAWWGHLEIVEVLLKRGADVSAADLHGQSPLHLAAMVGHLEIVEVLLKYGADVNAKDTMGATPLHLAARSGHLEIVEELLKNGADMNAQDKFGKTTFDISTDNGNEDLAEILQKLASDYKDDDDKGDYKDDDDKGDYKDDDDK

### K27-FKBP12^F36V^-3×FLAG tag

MDLGKKLLEAARAGQDDEVRILMANGADVNAHDTFGFTPLHLAALYGHLEIVEVLLKNGADVNADDSYGRTPLHLAAMRGHLEIVEVLLKYGADVNAADEEGRTPLHLAAKRGHLEIVEVLLKNGADVNAQDKFGKTAFDISIDNGNEDLAEILQKLGSGSGSMGVQVETISPGDGRTFPKRGQTCVVHYTGMLEDGKKVDSSRDRNKPFKFMLGKQEVIRGWEEGVAQMSVGQRAKLTISPDYAYGATGHPGIIPPHATLVFDVELLKLEASDYKDDDDKGDYKDDDDKG DYKDDDDK

### FKBP12^F36V^-K27-3×FLAG tag

MGVQVETISPGDGRTFPKRGQTCVVHYTGMLEDGKKVDSSRDRNKPFKFMLGKQEVIRGWEEGVAQMSVGQRAKLTISPDYAYGATGHPGIIPPHATLVFDVELLKLEGSGSGSGSDLGKKLLEAARAGQDDEVRILMANGADVNAHDTFGFTPLHLAALYGHLEIVEVLLKNGADVNADDSYGRTPLHLAAMRGHLEIVEVLLKYGADVNAADEEGRTPLHLAAKRGHLEIVEVLLKNGADVNAQDKFGKTAFDISIDNGNEDLAEILQKLASDYKDDDDKGDYKDDDDK GDYKDDDDK

### NbALFA-FKBP12^F36V^-3×FLAG

MEVQLQESGGGLVQPGGSLRLSCTASGVTISALNAMAMGWYRQAPGERRVMVAAVSERGNAMYRESVQGRFTVTRDFTNKMVSLQMDNLKPEDTAVYYCHVLEDRVDSFHDYWGQGTQVTVSSGSGSGSMGVQVETISPGDGRTFPKRGQTCVVHYTGMLEDGKKVDSSRDRNKPFKFMLGKQEVIRGWEEGVAQMSVGQRAKLTISPDYAYGATGHPGIIPPHATLVFDVELLKLEASDYKDDDDKGDYKDDDDKGDYKDDDDK

### Nb139-FKBP12^F36V^-3×FLAG

AQVQLQESGGGLVQAGGSLRLSCAASERTFSTYAMGWFRQAPGREREFLAQINWSGTTTYYAESVKDRTTISRDNAKNTVYLEMNNLNADDTGIYFCAAHPQRGWGSTLGWTYWGQGTQVTVSSGSGSGSMGVQVETISPGDGRTFPKRGQTCVVHYTGMLEDGKKVDSSRDRNKPFKFMLGKQEVIRGWEEGVAQMSVGQRAKLTISPDYAYGATGHPGIIPPHATLVFDVELLKLEASDYKDDDDKGDYKDDDDKGDYKDDDDK

### C10-FKBP12^F36V^-3×FLAG

MGSDLGKKLLEAAWHGQDDEVRILMANGADVNATDQSGMTPLHLAAWRGHLEIVEVLLKTGADVNAIDRWGKTPLHLAARIGHLEIVEVLLKAGADVNAQDKFGKTPFDLAIDNGNEDIAEVLQKAAGSGSGSMGVQVETISPGDGRTFPKRGQTCVVHYTGMLEDGKKVDSSRDRNKPFKFMLGKQEVIRGWEEGVAQMSVGQRAKLTISPDYAYGATGHPGIIPPHATLVFDVELLKLEASDYKDDDDKGDYKDDDDKGDYKDDDDK

---

**Figure S1.**
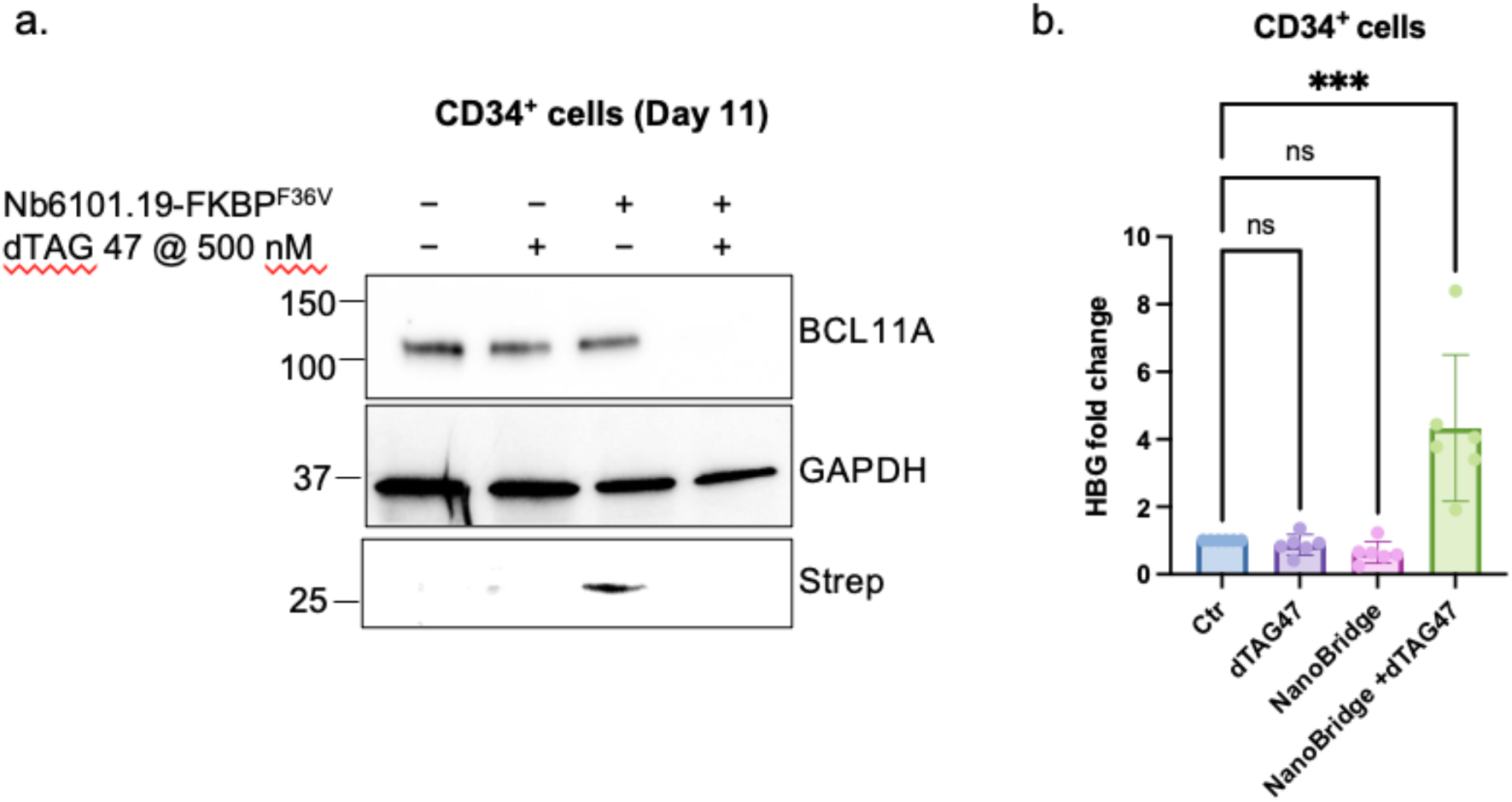
BCL11A degradation via the NanoBridge. (a) Immunoblots revealing that BCL11A degraded in CD34+ cells only in the presence of both the NanoBridge (Nb6101.19-FKBPF36V) and dTAG-47. (b) Fold change of HBG transcripts in CD34+ cells treated as indicated. CD34+ cells were collected from Day 13 and analyzed by RT-qPCR (mean ± s.d., n = 3, ***P < 0.001).

**Figure S2.**
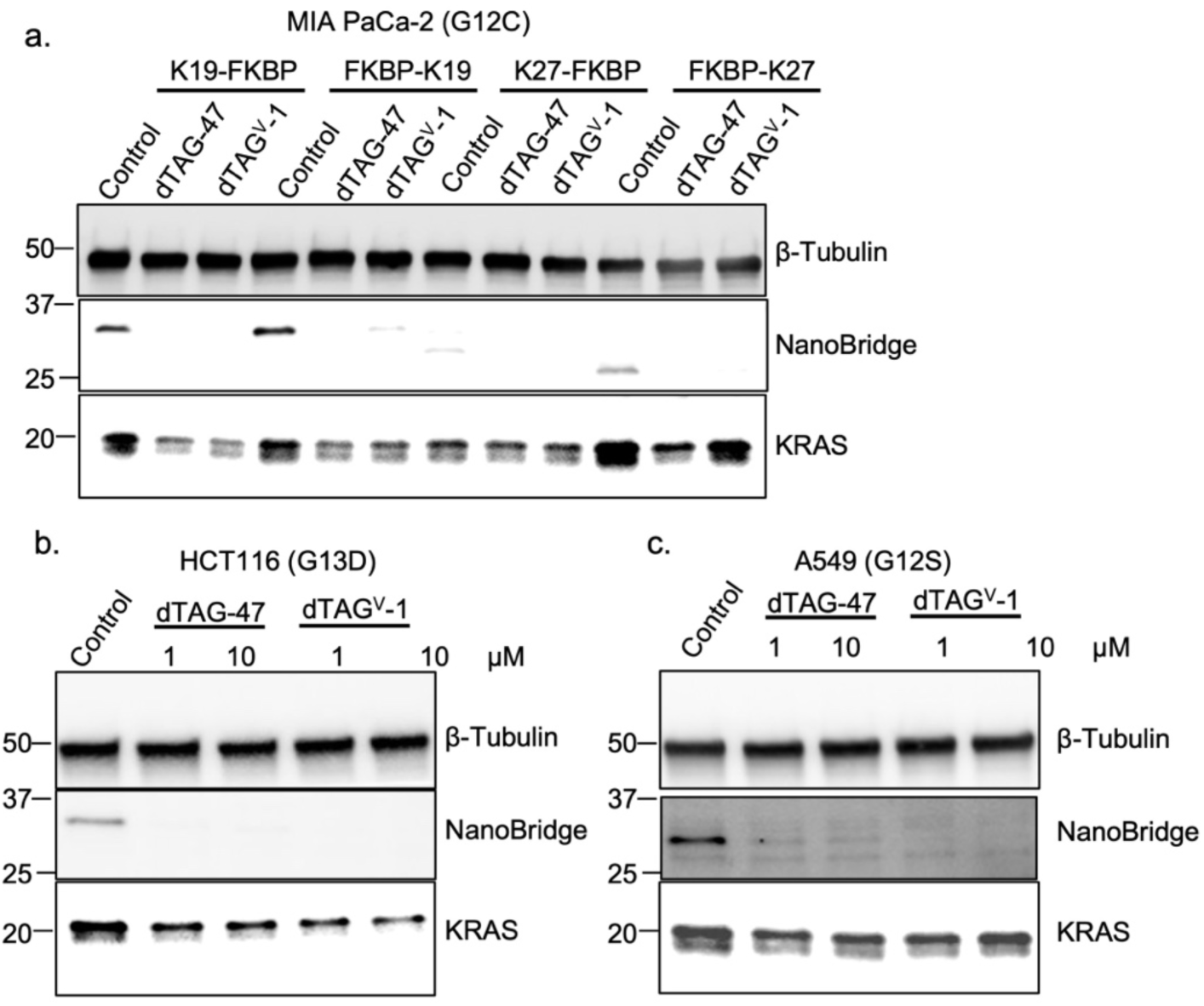
NanoBridge-induced targeted degradation of different KRAS mutants in cancer cells. Immunoblots revealing (a) degradation of KRAS (G12C) in Mia-PaCa 2 expressing different NanoBridge constructs upon treatment with 1 μM dATG47 or dTAGV-1 for 24h, (b) degradation of KRAS (G13D) in HCT116 cells and (c) degradation of KRAS(G12S) in A549 cells upon treatment with 1μM or 10 μM dATG47 or dTAGV-1 for 24h.

**Figure S3.**
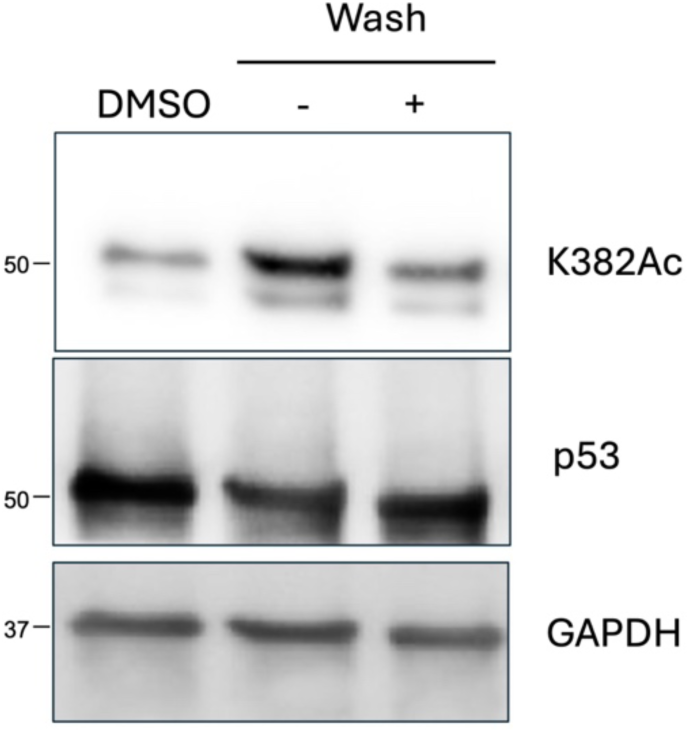
Reversibility of PTM induction. Immunoblots revealing removal of AceTAG-1 via a washout resulted in the reduction of p53 acetylation at K382 in 12 h. H1299 cells expressing Nb139-FKBP12K36V were treated with 500 nM AceTAG-1 for 3 h, and cells were washed with PBS for 3 times, and cultured in fresh complete medium for another 12 h. The cells not washed were treated with 500 nM AceTAG-1 for 15 h.

**Figure S4.**
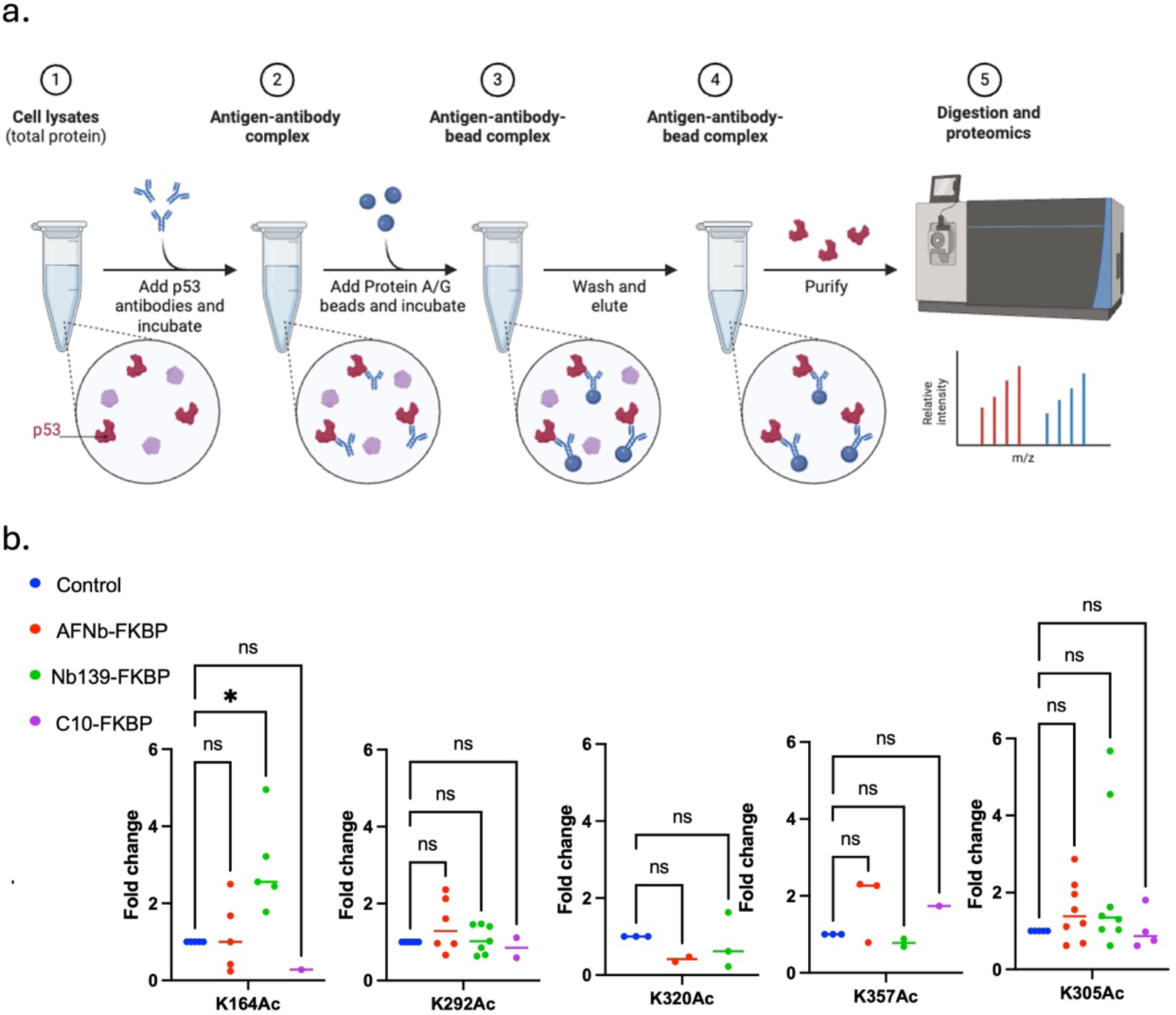
Proteomics to determine the fold change of p53 acetylated lysines in H1299 cells expressing different NanoBridge constructs treated with AceTAG-1. (a) Schematic description of the experimental workflow. Cells were treated with DMSO or AceTAG-1 (500 nM) for 16h and lysed; p53 was immunoprecipitated, digested and subjected to LC/MS/MS for analysis. (b) Fold change determined via mass spectrometry. Data represent a median of *n* = 4 biological replicates for AceTAG-1-treated cells. Note that peptides may not have been detected in every replicate but were detected and quantified in a minimum of two biological replicates to be included in this analysis. No statistically significant acetylation was detected for K292, K320, K357, and K305. Statistical significance was assessed via one-way ANOVA comparing DMSO-treated sample to AceTAG-1 treated samples. \**p* < 0.05; \*\*\**p* < 0.001; \*\*\*\**p* < 0.0001

**Table S1.**
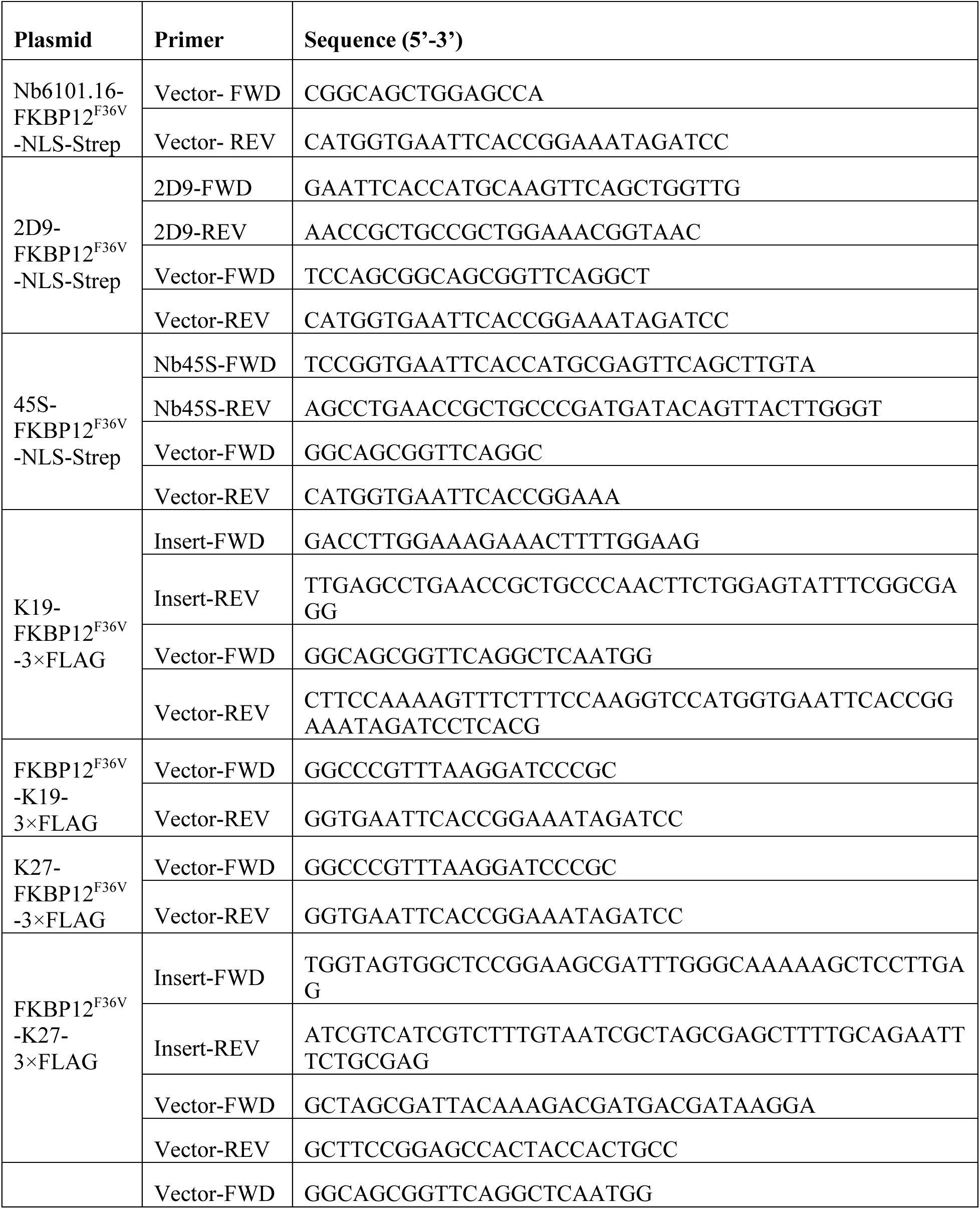
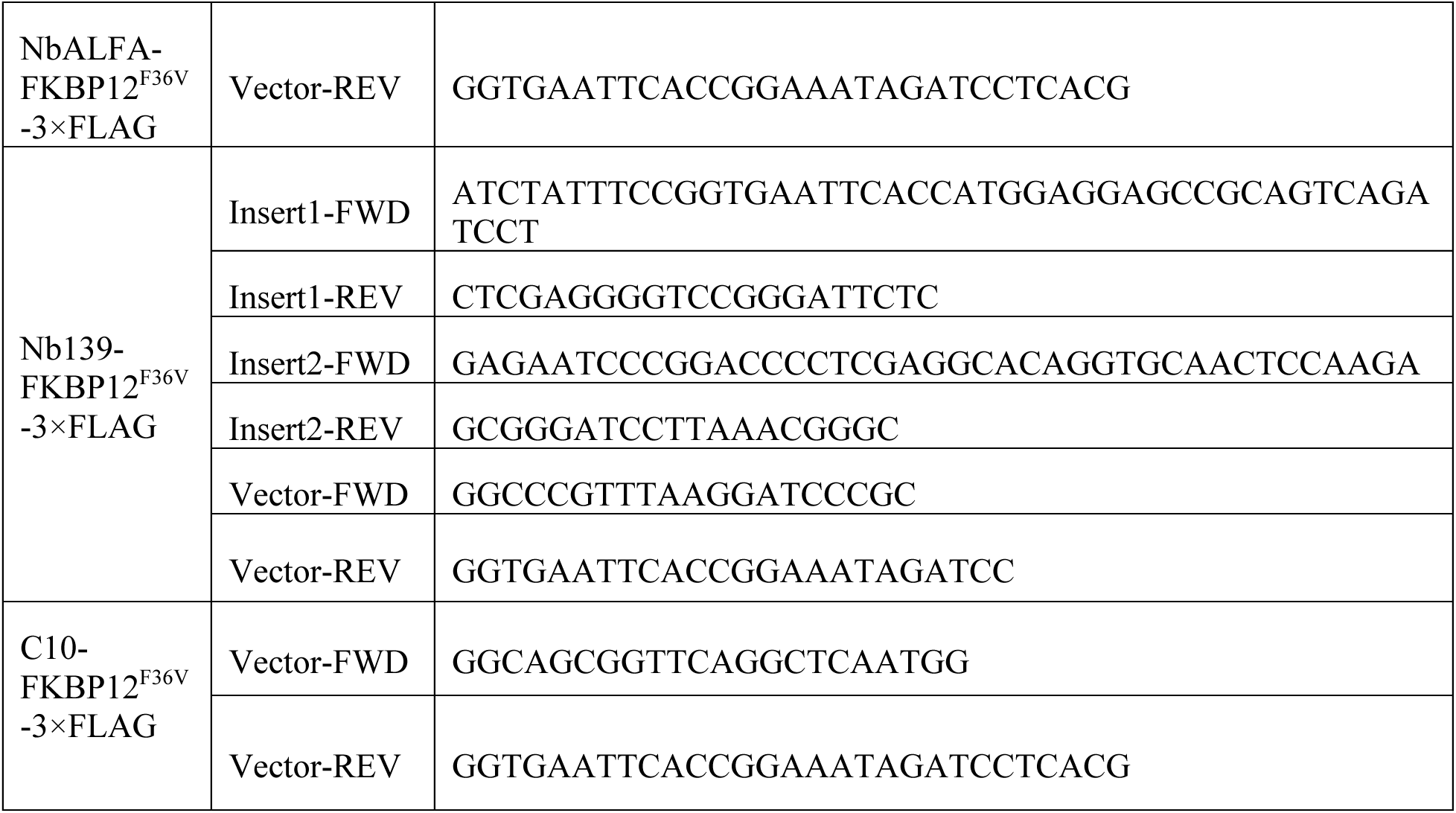
Primers used for plasmid construction.

**Table S2.**
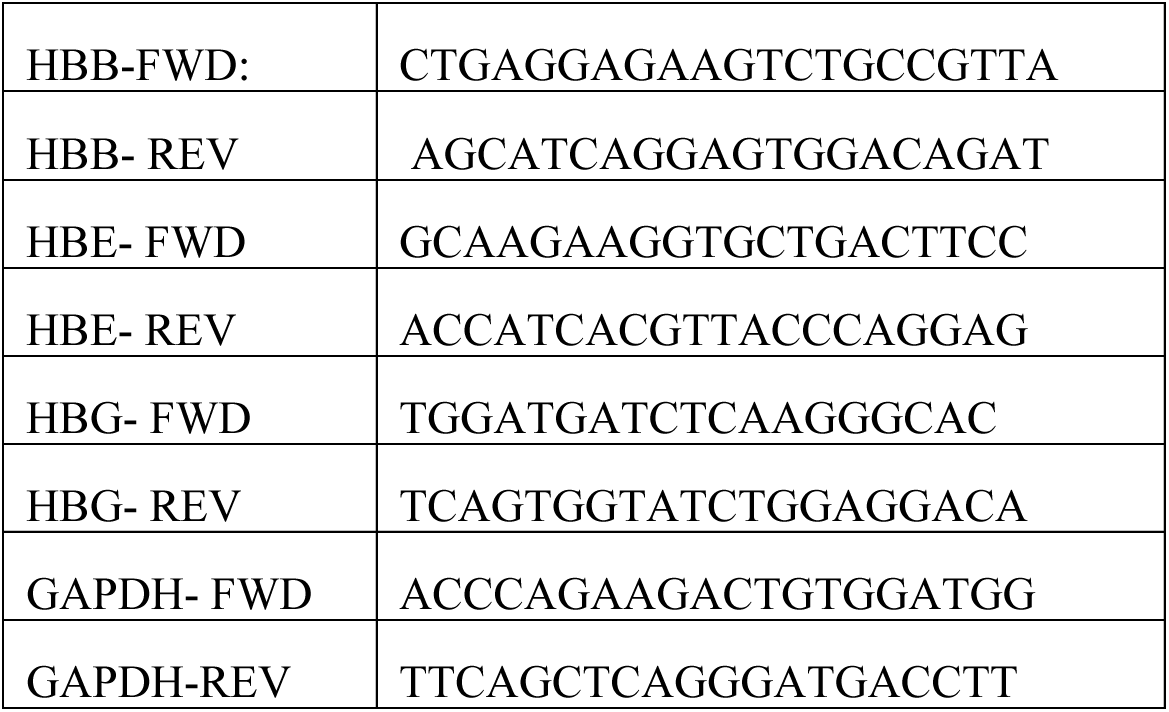
Primes used for RT-qPCR.

**Table S3.** C_q_ values of hemoglobin transcripts in CD34^+^ cells.

| C<sub>q</sub> values of hemoglobin transcripts in CD34<sup>+</sup> cells expressing Nb6101.19-FKBP12<sup>F36V</sup> |  |  |  |  |  |  |  |  |  |  |  |  |
| --- | --- | --- | --- | --- | --- | --- | --- | --- | --- | --- | --- | --- |
| CD34<sup>+</sup> cells (Day 13) |  |  |  |  |  |  |  |  |  |  |  |  |
|  | Untreated |  |  |  |  |  | Nb6101.19-FKBP12<sup>F36V</sup> |  |  |  |  |  |
|  | Cq 1 | Cq 2 | Cq 3 | Cq Avg | DCq | 2<sup>-DCq</sup> | Cq 1 | Cq 2 | Cq 3 | Cq Avg | DCq | 2<sup>-DCq</sup> |
| HBB | 23.09 | 23.03 | 22.95 | 23.02 | -0.43 | 1.35 | 26.08 | 25.69 | 26.03 | 25.93 | 1.88 | 0.27 |
| HBG | 23.31 | 23.06 | 23.28 | 23.21 | -0.23 | 1.17 | 24.07 | 24.06 | 24.47 | 24.2 | 0.15 | 0.90 |
| GAP<br>DH | 23.76 | 23.3 | 23.28 | 23.45 | / | / | 24.27 | 23.88 | 24.01 | 24.05 | / | / |
| C<sub>q</sub> values of hemoglobin transcripts in CD34<sup>+</sup> cells expressing different NanoBridge constructs |  |  |  |  |  |  |  |  |  |  |  |  |
| CD34<sup>+</sup> cells (Day 13) |  |  |  |  |  |  |  |  |  |  |  |  |
|  | Control |  |  |  |  |  | dATG47 |  |  |  |  |  |
|  | Cq 1 | Cq 2 | Cq 3 | Cq Avg | DCq | 2<sup>-DCq</sup> | Cq 1 | Cq 2 | Cq3 | Cq Avg | DCq | 2<sup>-DCq</sup> |
| HBB | 17.58 | 17.50 | 17.84 | 17.64 | -6.93 | 122.22 | 17.24 | 17.08 | 17.08 | 17.13 | -7.40 | 169.29 |
| HBG | 20.64 | 20.85 | 20.88 | 20.79 | -3.78 | 13.77 | 20.63 | 20.85 | 20.88 | 20.79 | -3.75 | 13.45 |
| GAP<br>DH | 24.50 | 24.37 | 24.85 | 24.57 | / | / | 24.39 | 24.44 | 24.78 | 24.54 | / | / |
|  | NbGFP-FKBP12<sup>F36V</sup> |  |  |  |  |  | Nb6101.19-FKBP12<sup>F36V</sup> |  |  |  |  |  |
|  | Cq 1 | Cq 2 | Cq 3 | Cq Avg | DCq | 2<sup>-DCq</sup> | Cq 1 | Cq 2 | Cq3 | Cq Avg | DCq | 2<sup>-DCq</sup> |
| HBB | 19.16 | 18.97 | 19.00 | 19.04 | -6.37 | 82.71 | 19.61 | 19.33 | 19.86 | 19.6 | -6.10 | 68.44 |
| HBG | 22.38 | 22.36 | 22.58 | 22.44 | -2.97 | 7.85 | 21.68 | 21.78 | 22.04 | 21.83 | -3.86 | 14.55 |
| GAP<br>DH | 25.17 | 25.02 | 26.05 | 25.41 | / | / | 25.72 | 25.66 | 25.71 | 25.70 | / | / |
|  | Nb2D9-FKBP12<sup>F36V</sup> |  |  |  |  |  | Nb45S-FKBP12<sup>F36V</sup> |  |  |  |  |  |
|  | Cq 1 | Cq 2 | Cq 3 | Cq Avg | DCq | 2<sup>-DCq</sup> | Cq 1 | Cq 2 | Cq3 | Cq Avg | DCq | 2<sup>-DCq</sup> |
| HBB | 19.12 | 18.51 | 18.81 | 18.81 | -6.79 | 110.40 | 19.46 | 19.27 | 19.37 | 19.37 | -5.95 | 61.82 |
| HBG | 21.34 | 21.17 | 21.40 | 21.30 | -4.30 | 19.65 | 21.31 | 21.51 | 21.69 | 21.50 | -3.81 | 14.06 |
| GAP<br>DH | 25.50 | 25.46 | 25.84 | 25.6 | / | / | 25.29 | 25.19 | 25.47 | 25.32 | / | / |

## Notes

### Competing Interest Statement

The authors have declared no competing interest.

